# Antibiotic Persistence Emerges from Cell-State-Driven Transcriptional Reprogramming

**DOI:** 10.64898/2026.02.16.705191

**Authors:** Qiang Liu, Zehui Yu, Xiaoli Liu, Amy Iverson, Tyler Simmons, Jason W. Rosch, Peijun Ma

## Abstract

Antibiotic persistence allows a subpopulation of bacterial cells to survive antibiotic treatment without acquiring resistance mutations, often contributing to treatment failure. Although many genes have been linked to persistence, their deletion rarely abolishes the phenotype, highlighting redundancy in persistence mechanisms. Moreover, the transient and heterogeneous nature of persister formation has made it difficult to resolve its molecular basis using bulk analyses. Here, we use bacterial single-cell RNA sequencing and functional assays in *Klebsiella pneumoniae* to demonstrate how transcriptional heterogeneity and redundancy in stress responses shape persistence outcomes. Using growth phase as a biologically meaningful axis of transcriptional variation, we reveal that even within an isogenic population, distinct transcriptional responses can be induced and co-contribute to survival. These responses are shaped by the cell’s pre-treatment transcriptional state and the mechanism of antibiotic action. Genetic and environmental perturbations, such as *rpoS* deletion and nutrient supplementation, shift pre-treatment cell states and alter persistence frequencies. Our findings establish the biological significance of transcriptional heterogeneity shaped by pre-treatment cell states, providing a systems-level framework for understanding persistence and suggesting strategies to enhance antibiotic efficacy by modulating cell states.

## Introduction

Even within genetically identical populations, bacterial cells can differ in their physiological states. This phenotypic heterogeneity, whether pre-existing or stress-induced, is widely viewed as a bet-hedging strategy to ensure population survival under stress^1–3^. A prominent example is antibiotic persistence, where a small subpopulation of cells survives otherwise lethal antibiotic treatment, contributing to treatment failure, recurrent infections, and the evolution of resistance^4–8^.

Antibiotic persisters can arise either as pre-existing cells that formed spontaneously or in response to other stresses, or as being induced upon antibiotic exposure^6,9,10^. For antibiotic-induced persisters, two steps are involved: first, the formation of persisters via transcriptional reprogramming in response to antibiotic stress; and second, the survival of persisters, often enabled by traits developed after the formation stage, such as metabolic dormancy, that help cells evade killing^9^. Thus, the trajectory of reprogramming induced upon antibiotic exposure is critical, as only certain directions lead to survival. However, how this direction is determined, how it unfolds across individual cells, and what factors govern it remain poorly understood.

This knowledge gap arises in part because the formation of persisters is a transient and dynamic process embedded within a phenotypically heterogeneous population. Bulk-level analysis often masks this heterogeneity, making it difficult to resolve the molecular mechanisms underlying persister formation^9^. For instance, previous studies have identified genetic pathways such as toxin-antitoxin systems, but deletion of these genes rarely eliminates persistence^11,12^, suggesting the mechanisms involved are redundant, conditional, or embedded within broader regulatory networks. Stress response pathways, including the stringent response, the general stress response (RpoS), and the SOS response, have also been proposed as a general route to antibiotic persistence, pointing to a role for transcriptional reprogramming in survival^12–19^. However, these findings are based largely on bulk data, leaving open the critical question of whether persistence arises through a single conserved transcriptional program or through multiple, divergent trajectories shaped by phenotypic heterogeneity. A central challenge is determining whether all cells follow a shared response to a given stress, or whether distinct subpopulations deploy parallel strategies to survive. Addressing this requires single-cell resolution to map the diversity of transcriptional responses underlying persistence^20^.

The advent of bacterial scRNA-seq has enabled genome-wide profiling of transcriptional heterogeneity at single-cell resolution^21–26^, offering new opportunities to dissect the dynamics of persister formation. A recent study applied this approach to identify a convergent persister-specific transcriptional state in *Escherichia coli*, but focused on pre-existing persisters formed before antibiotic exposure^27^. In contrast, understanding how antibiotics induce persistence, and whether different cell states follow distinct reprogramming paths under drug stress, requires systematic single-cell profiling during treatment.

Inspired by our recent discovery that meropenem, a β-lactam antibiotic that inhibits cell-wall synthesis, induces heterogeneous molecular responses including a subpopulation enriched for persisters^23^, we hypothesized that persister formation can be initiated through diverse mechanisms simultaneously triggered by antibiotics. Since antibiotic persistence varies across bacterial growth phases, with stationary-phase cells being the most tolerant^28,29^, we further hypothesized that cells at different growth phases exhibit distinct antibiotic responses that drive different levels of persistence.

To test these hypotheses, we focused on *Klebsiella pneumoniae*, a clinically important pathogen associated with recurrent and multidrug-resistant infections^30^. We used scRNA-seq to profile its transcriptional responses to antibiotics across lag, exponential, and stationary phases, leveraging growth phase as a biologically meaningful axis of variation in pre-treatment cell states. Our findings reveal that antibiotic survival can be achieved through divergent transcriptional reprogramming pathways within the same isogenic population, shaped by both the cell’s pre-treatment transcriptional state and the antibiotic’s mechanism of action. Moreover, we show that altering the pre-treatment cell states, through genetic or nutrient perturbation, can modulate persistence levels.

Together, these results demonstrate how pre-treatment cell states shape transcriptional reprogramming and survival during antibiotic exposure, providing direct evidence for the biological significance of transcriptional heterogeneity in antibiotic survival and persistence. This work establishes: (i) a systems-level framework that provides mechanistic insight into why persistence cannot be eliminated by targeting single genes; (ii) a potential strategy to enhance antibiotic efficacy by modulating bacterial cell states; and (iii) a foundation for broader single-cell profiling across bacterial species and for *in vivo* characterization of heterogeneity and drug responses.

## Results

### Profiling pre-treatment cell states at various growth phases that are linked to different levels of persistence

To profile transcriptional reprogramming across bacterial growth phases, we first mapped the “pre-treatment cell states” using scRNA-seq in cultures prior to antibiotic treatment. We used an optimized version of BacDrop protocol (BacDrop_V2) with improved coverage (see Methods and Supplementary Fig. 1). We define “cell state” as a transcriptomic profile representing a cell’s regulatory and physiological status, serving as a proxy for its proteomic and metabolic activity. Although transcriptomes do not fully capture cellular function, they provide a predictive snapshot of pathways involved in stress adaptation and survival. While bacterial cultures are traditionally categorized by bulk growth phases, it remains unclear whether these phases represent uniform physiological states or heterogeneous mixtures of distinct cell states. This distinction is critical, as population-level measurements obscure the single-cell heterogeneity that may shape antibiotic response and survival.

To define bacterial cell states at single-cell resolution before antibiotic exposure, we performed scRNA-seq on a clinical isolate of *K. pneumoniae* MGH66 cultures at early lag (10 minutes post-transfer to fresh medium, LAG10), exponential (EXP), and stationary phases (STA) under unperturbed conditions (Fig. 1a). To minimize carryover from the inoculum, cultures were serially diluted three times before sampling EXP cells (∼2 hours after dilution into fresh medium). Transcriptional cell states were identified using unsupervised clustering followed by pathway enrichment analysis, enabling functional annotation of each cluster (Fig. 1b,c, Supplementary Fig. 2). While we refer to these as discrete “states,” they likely lie along a continuum and represent overlapping transcriptional programs dynamically weighted over time. To ensure robust annotation and reproducibility, we included two biological replicates per condition and excluded clusters found in only one replicate.

**Figure 1.**
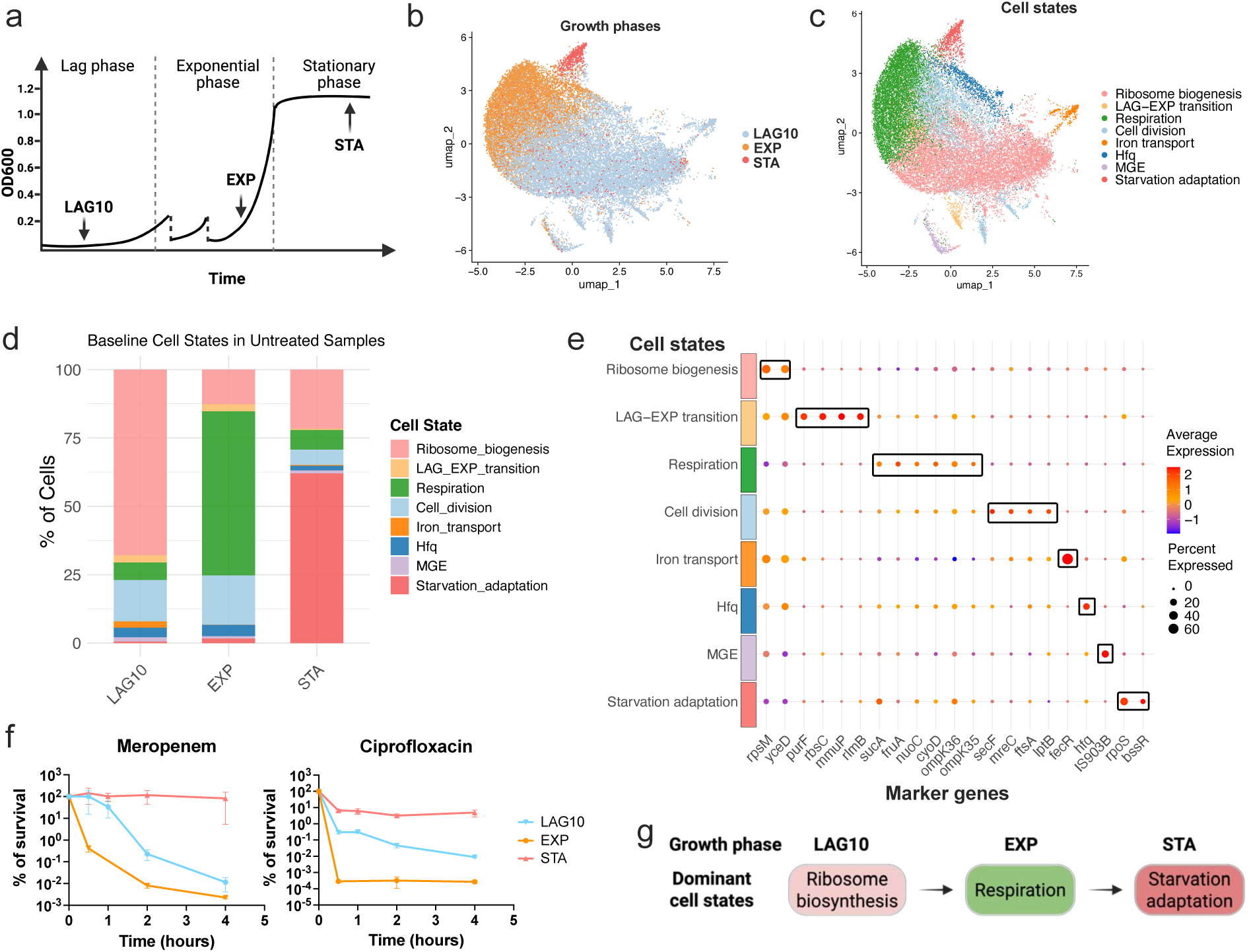
Growth-phase-specific transcriptional cell states underlie antibiotic persistence. **a,** Schematic of the sampling strategy for scRNA-seq across three growth phases: early lag (10 minutes post-transfer to fresh medium, LAG10), exponential (EXP), and stationary phases (STA) under unperturbed conditions. **b,** UMAP projection of single-cell transcriptomes from LAG10, EXP, and STA samples, colored by growth phase. Two independent biological replicates (n = 2) were included for each condition. **c,** UMAP embedding colored by eight annotated cell states, identified based on gene expression markers across all growth phases. **d,** Distribution of cell states across growth phases. Cell state proportions were averaged from two biological replicates. **e,** Selected marker genes representative of each annotated cell state. **f,** Killing dynamics of *K. pneumoniae* MGH66 following treatment with meropenem (2 µg/ml) or ciprofloxacin (2.5 µg/ml), initiated at different growth phases. Persistence levels differed by growth phase. CFU was quantified over time; data represent three independent biological replicates (n = 3), and error bars denote standard deviation. **g,** Summary of growth-phase-dominated cell states.

We identified eight reproducible transcriptional cell states across three growth phases, with each phase characterized by a dominant expression profile (Fig. 1c–e, Supplementary Tables 1–3). At the LAG10 phase, ∼70% of cells exhibited a ribosome biogenesis program. During the EXP phase, ∼60% of cells exhibited relative enrichment of respiration-associated transcripts compared to other states, while ∼10% retained a ribosome-associated transcriptional profile. By the STA phase (16 hours post-dilution), ∼60–70% of cells adopted a starvation-adaptation state, marked by *rpoS* and biofilm gene upregulation. These findings reveal structured heterogeneity within bacterial growth phases, with transcriptional profiles correlating with differential antibiotic survival: at the population level, stationary-phase cells showed the highest persistence, while exponential-phase cells were the most susceptible (Fig. 1f). Of note, to maintain analytical consistency, we applied the same minimum gene-detection cutoff across all three growth phases. However, stationary-phase bacteria are known to yield substantially fewer detectable transcripts per cell, a biological property reported in multiple bacterial single-cell studies, including *E. coli* ^21,27^ and *Bacillus subtilis*^22^. As a result, fewer stationary-phase cells passed this uniform filtering threshold (Supplementary Table 1). To assess whether this difference affected state identification, we performed an additional analysis in which the minimum gene-detection cutoff was specifically relaxed for stationary-phase cells to achieve comparable cell numbers to the LAG10 and EXP phases. This analysis recovered the same major transcriptional states and overall structure, yielding results consistent with those obtained using a uniform cutoff (Supplementary Fig. 3; Supplementary Table 1). We note that despite this validation, stationary-phase sampling depth remains lower than that of lag and exponential phases, which may limit detection of rare transcriptional subpopulations.

These results demonstrate that bacterial growth phases comprise transcriptionally distinct cell states that are associated with functional differences in antibiotic survival. These transcriptional states capture dominant transcriptional programs that are differentially enriched across growth phases, reflecting shifts in physiological emphasis rather than mutually exclusive or sequential activation of individual processes (Fig. 1g). Notably, a “cell division” state was identified across all three growth phases (Fig. 1c,d), marked by core division genes such as *ftsA* (Fig. 1e). However, closer inspection revealed that cells within this state exhibited distinct marker profiles associated with ribosome biosynthesis or respiration, reflecting their respective growth phases. In subsequent experiments, we use these growth phase-defined populations, each with a characteristic distribution of cell states, as a biologically meaningful axis to test how pre-treatment cell states influence transcriptional reprogramming and persistence under antibiotic exposure.

### Meropenem induced heterogeneous transcriptional reprogramming across growth phases

We next profiled the transcriptional reprogramming induced by meropenem across bacterial growth phases characterized by distinct cell state structures. Using the same sampling scheme as in Fig. 1a, we exposed *K. pneumoniae* cultures from LAG10, EXP, and STA phases to meropenem at a lethal concentration (40× of minimal inhibitory concentration (MIC)). For the LAG10 and EXP cells, we collected samples when ∼40–60% of cells had been killed based on quantification of colony forming unit (CFU) on LB agar plates. For STA cells that showed high-level antibiotic tolerance (Fig. 1f), we collected cells at 15 minutes after antibiotic exposure.

Upon meropenem exposure, scRNA-seq results revealed that three new cell states emerged: 1) the heat shock response that is marked by upregulation of *dnaK*, *groES*, and *groEL*; 2) *osmB* upregulation accompanied with induced expression of two small RNAs *csrB* and *csrC*; 3) and a cell state with both iron transport and heat shock upregulation (Fig. 2a–c, Supplementary Fig. 4, Supplementary Tables 4, 5). Across all three states, we observed induction of *lpp* (annotated as *lpp1* in *K. pneumoniae* MGH66 genome), which encodes Braun’s lipoprotein, the major outer membrane protein that covalently links the outer membrane to the peptidoglycan layer^31^, suggesting a shared response involving envelope stabilization under stress (Fig. 2c).

**Figure 2.**
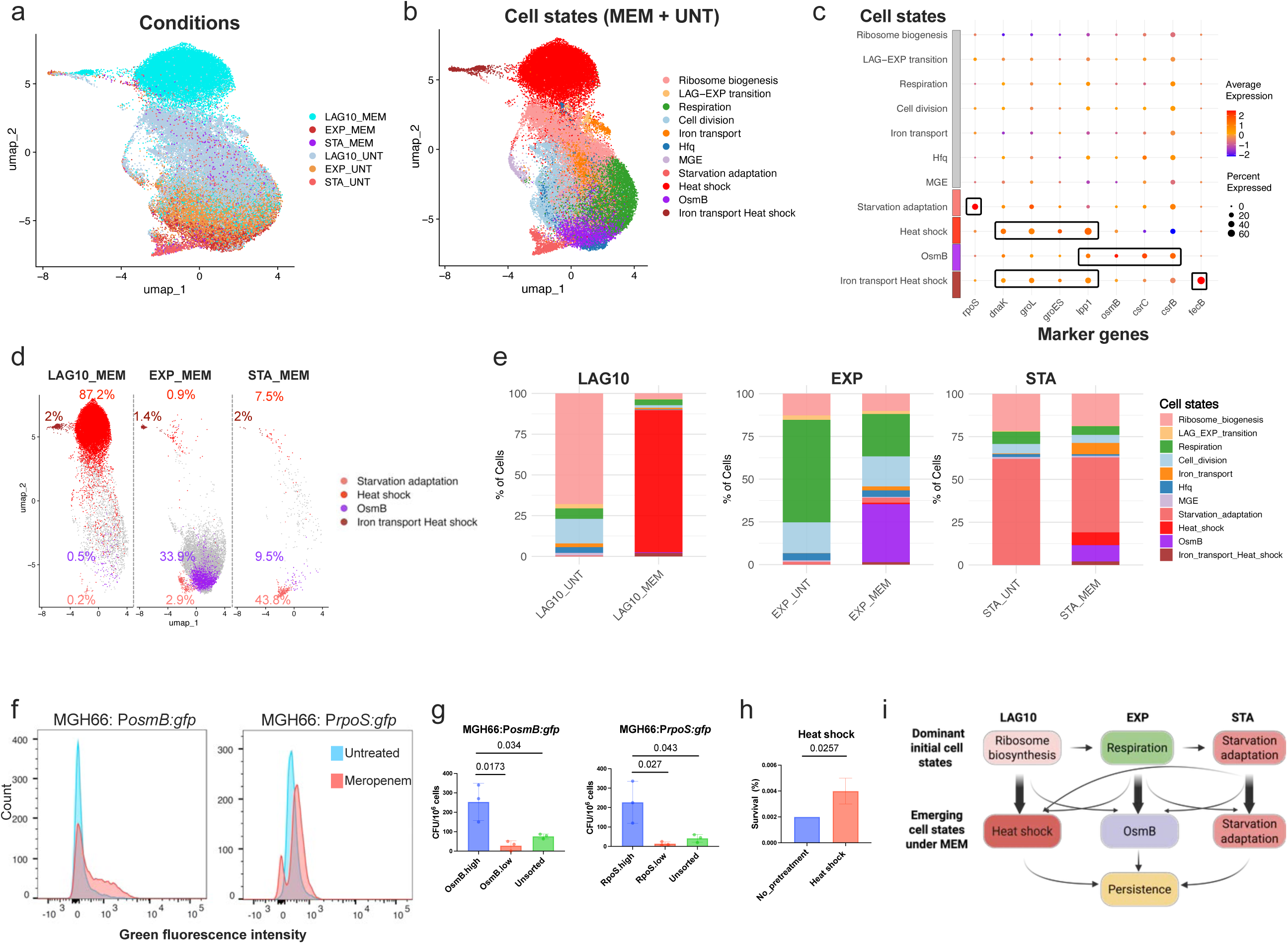
Growth-phase-specific transcriptional reprogramming under meropenem reveals divergent stress responses co-contributing to antibiotic survival. **a,** UMAP embedding of single-cell transcriptomes from six conditions: three untreated (UNT) samples at early lag (10 minutes post-transfer to fresh medium, LAG10), exponential (EXP), and stationary phases (STA) and three meropenem-treated (MEM) samples collected from each corresponding growth phase (LAG10_MEM, EXP_MEM, and STA_MEM). Each condition includes two independent biological replicates (n = 2). **b,** Same UMAP as in **a**, colored by transcriptional cell states. In addition to the eight shared states identified in untreated cells, three new states induced by meropenem treatment were identified: Heat shock, OsmB, and Iron transport + Heat shock. **c,** Dot plot showing representative marker genes for the three meropenem-induced cell states and the Starvation adaptation cell state. **d,** UMAPs of meropenem-treated cells alone, separated by growth phase. The percentage of cells in each state is labeled to highlight dominant and minor transcriptional responses (Pink, Starvation adaptation; red, Heat shock; purple, OsmB; brown, Iron transport + Heat shock) **e,** Bar plot showing the distribution of all cell states, including the eight original states and three new states, across meropenem-treated samples. Percentages represent the average from two biological replicates. **f,** Flow cytometry validation of scRNA-seq-inferred heterogeneity using GFP reporter strains. Promoters of *osmB* or *rpoS* were used to drive GFP expression in *K. pneumoniae* MGH66. Variable fluorescence intensity confirms heterogeneous expression across the population. This experiment was performed using three independent biological replicates (n = 3), and the data from one representative replicate is shown. **g,** Functional validation of marker genes using fluorescence-activated cell sorting (FACS). EXP-phase cells with high GFP expression (top 10%) for *osmB* (OsmB.high) or *rpoS* (RpoS.high) were sorted and compared to low-expression (bottom 10%) and unsorted populations. Following 4-hour meropenem treatment (2 µg/ml), high-expressing cells showed increased survival, indicating that *osmB* and *rpoS* contribute to cell survival. Three independent biological replicates were performed (n =3). Bar plots show means of three biological replicates, and error bars represent standard deviation. Student’s t-test was used for the statistical analysis *p*-values are indicated on the plot. **h,** Heat shock preconditioning enhances survival under meropenem. EXP-phase cells were pre-treated at 42°C for 30 min prior to antibiotic exposure. Pre-treated cells exhibited higher survival compared to controls, supporting a protective role for heat shock priming. This experiment was performed with three biological replicates (n = 3). Bar plots show means ± standard deviation. Student’s t-test was used for the statistical analysis and *p*-values are indicated on the plot. **i,** Schematic model illustrating meropenem-induced persistence across growth phases. Each growth phase is characterized by a dominant pre-treatment cell state that guides distinct transcriptional responses to meropenem. These include Heat shock (with and without Iron transport), OsmB, and Starvation adaptation, which together enhance survival.

Although these three new cell states were observed in cells treated with meropenem across all growth phases, major transcriptional responses differed dramatically by growth phase (Fig. 2d,e). In LAG10 cells treated with meropenem, 79.6–94.7% of the population activated the heat shock response (Fig. 2d,e). Given that LAG10 cells were actively engaged in ribosome biogenesis (Fig. 2e), this response likely mitigates proteotoxic stress from misfolded or stalled translation products. In addition, 0.1–1.0% of LAG10 cells upregulated *osmB* and 2% of cells showed high-level expression of both iron transport genes and heat shock response.

EXP-phase cells also showed a heterogenous reprogramming under meropenem treatment, consistent with our previous findings^23^. Approximately 34% upregulated *osmB*, while smaller subpopulations activated starvation adaptation (2.7–3.1%), heat shock programs (0.8–1.0%), or both heat shock and iron transport responses (1.2–1.5%) (Fig. 2d,e; Supplementary Table 4). Of note, a starvation-adaptation population was also detected in untreated EXP-phase cells (1.6%) (Fig. 1d), suggesting that these cells may emerge spontaneously during growth and are subsequently enriched, or induced, under antibiotic stress. Given that EXP cultures were serially diluted prior to sampling (Fig. 1a), these cells are unlikely to represent carryover from the inoculum because only 0.2–0.8% of untreated LAG10-phase cells were in the starvation adaptation state (Fig. 1d, Supplementary Table 1).

STA cells showed high-level antibiotic tolerance and had been considered as dormant cells (Fig. 1f). However, unexpectedly, we observed that 8.9–10% of STA cells upregulated *osmB*, 6.8–8.1% of STA cells showed heat shock response, and 1.9–2.1% of STA cells showed both iron transport and heat shock responses under meropenem treatment (Fig. 2d,e; Supplementary Table 4). Notably, all of these responding cells upregulated *lpp1* (Fig. 2c). In addition, we also observed a significant reduction of cells in the starvation adaptation state and a 20-fold increase of cells in the iron transport state. Similarly to the untreated condition, stationary-phase cells treated with meropenem yielded fewer cells passing quality control when a uniform minimum gene-detection cutoff was applied. To assess whether this affected transcriptional state identification, we performed an additional analysis in which the minimum gene-detection threshold was specifically relaxed for STA cells. This analysis recovered transcriptional states and overall structure consistent with the primary analysis, indicating that the conclusions are robust to differences in cell number and detection depth (Supplementary Fig. 5; Supplementary Table 4). Experimentally, using a GFP reporter driven by the native *lpp1* promoter, we validated that STA cells indeed upregulated *lpp1* (Supplementary Fig. 6), confirming that meropenem triggers transcriptional reprogramming in a subset of STA cells under our experimental conditions.

Together, these results demonstrate that meropenem induces heterogeneous transcriptional responses, with profiles varying by growth phase. The dominant responses observed, such as heat shock in LAG10 cells and *osmB*-upregulation in EXP cells, highlight the influence of pre-treatment cell states on transcriptional reprogramming. LAG10, EXP, and STA populations differ not only in baseline physiology but also in how they reprogram under stress, underscoring the importance of growth-phase-specific cell states in shaping antibiotic responses.

### Antibiotic survival is enhanced from multiple mechanisms simultaneously induced by meropenem

The observation that multiple stress responses were induced motivated us to test the functional relevance of the observed transcriptional heterogeneity. Specifically, in EXP-phase cells treated with meropenem, three transcriptional responses, including *osmB* upregulation accompanied with upregulation of *csrB* and *csrC*, starvation adaptation marked by high-level expression of *rpoS*, and heat shock response, were identified as candidate survival-associated programs.

To assess their relationship to antibiotic survival, we measured survival rates using GFP reporter strains driven by the *osmB* or *rpoS* promoter. After meropenem exposure, EXP-phase cells were sorted into high- and low-GFP populations (top/bottom 10%), excluding dead cells using DAPI (Fig. 2f, Supplementary Fig. 7a–e). High expressers (OsmB.high or RpoS.high) exhibited a 4 to 10-fold increase in survival compared to low expressers and unsorted controls (Fig. 2g). To address the possibility that higher GFP signal reflects non-growing cells accumulating GFP non-specifically, we included a control reporter strain in which GFP is driven by the promoter of a housekeeping gene (*fusA*), whose expression is high and relatively uniform across the population with minimal cell-to-cell heterogeneity under our experimental conditions. Using this control, we show that sorting FusA.high and FusA.low cells does not result in a significant difference in survival, indicating that the survival advantage observed for stress-responsive reporters is not due to nonspecific GFP accumulation (Supplementary Fig. 7g,h).

To examine whether activation of the heat-shock response broadly modulates survival, we pre-treated EXP cells at 42°C for 30 minutes prior to antibiotic exposure. Heat-shocked EXP cells showed reproducible increase in survival following meropenem treatment (Fig. 2h), consistent with the protective role of heat shock observed in LAG10 cells. This pre-treatment likely primes cells to mitigate proteotoxic stress, potentially explaining the greater tolerance of LAG10 cells relative to untreated EXP cells (Fig. 1f).

Together, these results show that meropenem treatment induces divergent transcriptional responses that collectively modulate survival outcomes (Fig. 2i). This redundancy provides a potential explanation for why perturbation of individual genes often fails to eliminate surviving cells. While these experiments do not establish direct causal relationships between specific transcriptional states and persistence at the single-cell level, they demonstrate that transcriptional heterogeneity shapes antibiotic survival in a growth-phase-dependent and context-specific manner, setting the stage for the emergence of persistence under sustained antibiotic stress.

### Divergent transcriptional reprogramming induced across mechanistically distinct antibiotics

The findings under meropenem treatment raise another critical question: can divergent transcriptional reprogramming also be triggered by antibiotics with distinct cellular targets, such as those inhibiting DNA replication? To address this question, we treated *K. pneumoniae* cells from LAG10, EXP, and STA phases with a lethal concentration (80x MIC) of ciprofloxacin, a DNA synthesis-inhibiting antibiotic, and profiled their reprogramming by scRNA-seq. Ciprofloxacin and related fluoroquinolones are known to induce the SOS response^32^, centered on *recA* and downstream DNA repair genes.

Under ciprofloxacin treatment, unsupervised clustering revealed four new transcriptional cell states characterized by *recA* upregulation (Fig. 3a–c, Supplementary Fig. 8, Supplementary Tables 6,7). While all four clusters shared activation of the SOS response, they were transcriptionally distinct due to differences in co-regulated stress response programs (Fig. 3c). We named these clusters based on their predominant cell-of-origin: SOS_EXP and three SOS_LAG clusters. The SOS_EXP cluster co-upregulated canonical heat shock genes including *dnaK*, *groL*, and *groES*, in addition to *recA*. In contrast, SOS_LAG clusters upregulated *recA* and *recN*, a gene involved in recombination repair of DNA double stranded breaks, but did not activate the heat shock program at the time of sampling. Among the SOS_LAG clusters, the degree of *recN* upregulation varied, with SOS_LAG1 showing the highest expression (Fig. 3c).

**Figure 3.**
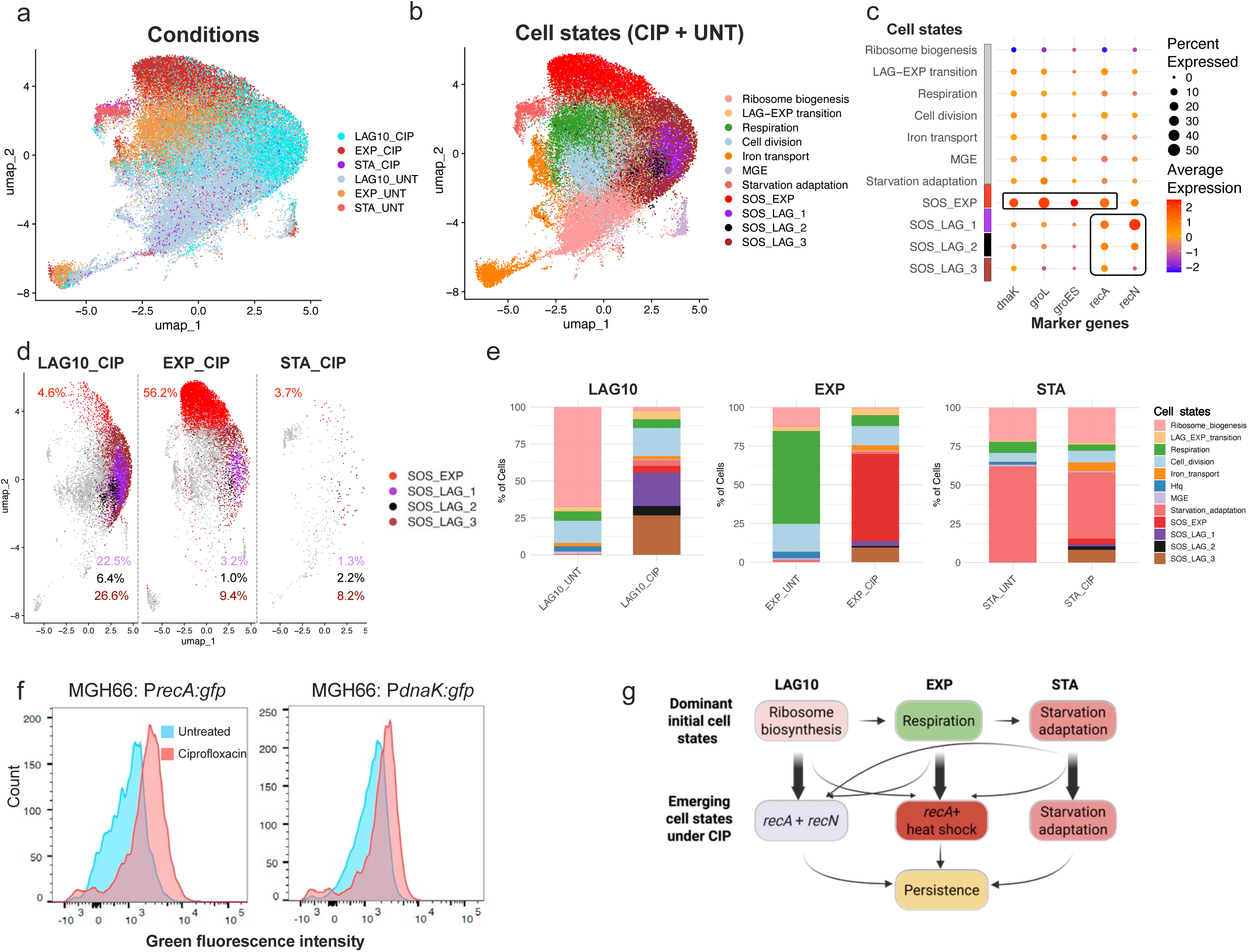
Divergent transcriptional reprogramming is also induced under ciprofloxacin treatment. **a,** UMAP embedding of single-cell transcriptomes from six conditions: three untreated (UNT) samples at early lag (10 minutes post-transfer to fresh medium, LAG10), exponential (EXP), and stationary phases (STA) and three ciprofloxacin-treated (CIP) samples collected from each corresponding growth phase (LAG10_CIP, EXP_CIP, and STA_CIP). Each condition includes two independent biological replicates (n = 2). **b,** Same UMAP as in **a**, colored by transcriptional cell states. In addition to the eight shared states identified in untreated cells, four new ciprofloxacin-induced states were identified: SOS_EXP, SOS_LAG_1, SOS_LAG_2, and SOS_LAG_3, labeled based on their growth-phase origin. **c,** Dot plot showing representative marker genes for the four ciprofloxacin-induced states. **d,** UMAPs of ciprofloxacin-treated cells alone, separated by growth phase. The percentage of cells in each state is labeled to highlight dominant and minor transcriptional responses. **e,** Bar plot showing the distribution of all cell states, including the eight shared states and four new states, across ciprofloxacin-treated samples. Percentages represent the average from two biological replicates. **f,** Flow cytometry validation of scRNA-seq-inferred heterogeneity using GFP reporter strains. Promoters of *recA* or *dnaK* were used to drive GFP expression in *K. pneumoniae* MGH66. Heterogeneous fluorescence signals confirm variable expression across the population. Shown is a representative plot from three independent biological replicates (n = 3). **g,** Schematic model illustrating ciprofloxacin-induced transcriptional reprogramming. Each growth phase is characterized by a dominant pre-treatment cell state, which shapes distinct stress responses upon ciprofloxacin exposure.

These SOS clusters were not exclusive to a single growth phase but appeared across all three (Fig. 3d,e, Supplementary Table 6). However, their relative abundance and transcriptional profiles were strongly phase dependent. In EXP-phase cells, approximately 53.1–59.3% co-induced the SOS and heat shock responses, and ∼12.6–14.4% upregulated *recA* and *recN* without activating heat shock genes. Flow cytometry using GFP reporters confirmed *recA* and heat shock gene induction in EXP cells, validating their co-upregulation at the EXP growth phase under ciprofloxacin treatment (Fig. 3f, Supplementary Fig. 9). In contrast, ∼55% of LAG10 cells upregulated *recA* and *recN* without heat shock gene induction (SOS_LAG). Similar to the response observed under meropenem, STA cells, traditionally regarded as dormant, displayed minimal but detectable transcriptional activity under ciprofloxacin treatment. Most responding cells exhibited *recA* activation together with weak *recN* upregulation within the SOS_LAG3 cluster. Similarly, for ciprofloxacin-treated samples, STA cells yielded fewer cells passing quality control under a uniform gene-detection cutoff; relaxing this threshold for STA cells recovered transcriptional states consistent with the primary analysis (Supplementary Fig. 10; Supplementary Table 6). Using a GFP reporter driven by the native *recA* promoter, we validated that a subset of STA cells indeed upregulated *recA* (Supplementary Fig. 11), confirming that ciprofloxacin also induces transcriptional reprogramming in these cells under our experimental conditions.

Together, these results demonstrate that while both LAG10 and EXP cells activate the SOS response to ciprofloxacin, the co-activation of additional stress programs diverges between phases (Fig. 3g). EXP cells prominently co-induce canonical heat shock genes, whereas LAG10 cells primarily co-induce the downstream DNA repair gene *recN*. This divergence may reflect phase-specific differences in reprogramming capacity, transcriptional burden, or stress thresholds that are rooted in their distinct pre-treatment physiological states: EXP cells are predominantly engaged in respiration, while LAG10 cells are characterized by active ribosome biosynthesis (Fig. 3g). While these data reveal clear phase-dependent heterogeneity within the SOS response, the functional consequences of this heterogeneity for survival under ciprofloxacin treatment were not directly tested and remain an open question.

Combined with our findings under meropenem treatment, these results support a general principle: antibiotics can induce divergent transcriptional reprogramming within and across growth phases, shaped not by drug mechanism alone, but by the interplay between a cell’s pre-treatment transcriptional state and the antibiotic’s mode of action. Crucially, this divergent reprogramming is functionally important for survival, as demonstrated in EXP cells under meropenem treatment (Fig. 2g,h), and likely applicable to ciprofloxacin-induced states as well. Notably, complementary flow cytometry experiments across a range of antibiotic concentrations indicate that the same stress-responsive transcriptional programs are induced under lower, more clinically relevant doses, although their timing and magnitude are concentration dependent (Supplementary Fig. 12). Our findings therefore provide a systems-level view of antibiotic survival at the early stage of persister formation, emphasizing how pre-treatment cell states shape transcriptional responses and survival, and pointing to new opportunities to predict or manipulate bacterial outcomes by modulating cell states.

### *RpoS* deficiency alters levels of antibiotic persistence via modulation of pre-treatment cell states

Building on our observations in wild-type *K. pneumoniae*, we hypothesized that antibiotic persistence levels could be altered by perturbing the pre-treatment cell states prior to drug exposure. To test this, we deleted *rpoS*, a conserved sigma factor that regulates general stress responses across bacterial species and has been implicated as a key gene in antibiotic persistence^12,13^.

Consistent with previous findings^18^, *rpoS* deletion significantly reduced persistence in STA-phase cells. Notably, the same deletion markedly increased survival during the LAG10 phase under both meropenem and ciprofloxacin treatment, while having no effect in EXP-phase cells (Fig. 4a). This pattern was reproduced in *E. coli* K-12, with *rpoS* deletion decreasing persistence in STA cells, increasing survival in LAG cells, and showing no effect in EXP cells (Supplementary Fig. 13), indicating that the phenomenon is not species-specific. Although enhanced persistence upon *rpoS* deletion has been documented in *E. coli* and *Pseudomonas aeruginosa*^16^, the mechanisms underlying these contrasting, growth phase-dependent outcomes have remained unclear.

**Figure 4.**
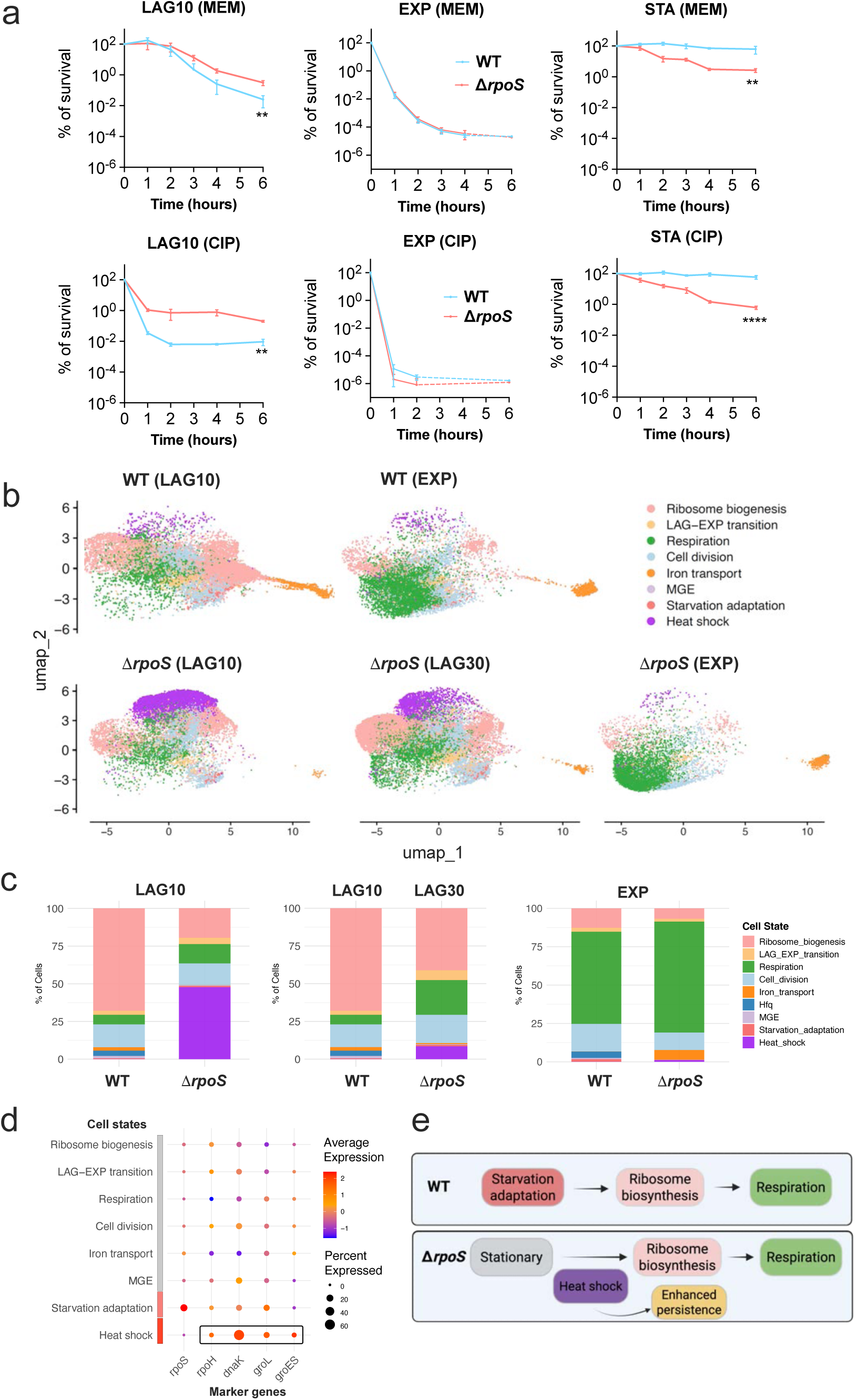
RpoS deficiency alters levels of antibiotic persistence via modulation of pre-treatment cell states. **a,** Antibiotic killing dynamics of wild-type (WT; blue) and Δ*rpoS* (red) strains of *K. pneumoniae* MGH66 following treatment with meropenem (MEM, top) or ciprofloxacin (CIP, bottom) at early lag (10 minutes post-transfer to fresh medium, LAG10), exponential (EXP), and stationary phases (STA). RpoS deletion reduced persistence at the STA phase, increased persistence at the LAG10 phase, and showed minimal effect at the EXP phase. Curves represent the mean of three independent biological replicates (n = 3); dashed lines indicate values under the limit of detection. Error bars indicate standard deviation. Student’s t-test was used for the statistical analysis to compare CFU from two strains at 6 hours after treatment. The *p*-values are 0.0083 (LAG10, MEM), 0.0032 (LAG10, CIP), 0.0078 (STA, MEM), and <0.0001 (STA, CIP). No significant difference was identified in EXP phase. **b,** UMAPs comparing single-cell transcriptional states of WT (top) and Δ*rpoS* (bottom) strains across LAG10, LAG30 (30 minutes post-transfer to fresh medium), and EXP phases. A distinct cluster of cells expressing heat shock genes emerged in the Δ*rpoS* strain at LAG10, decreased at LAG30, and was largely absent at EXP. Each condition includes two biological replicates (n = 2) except for Δ*rpoS* at EXP phase, which includes only one replicate due to a technical error during library construction. **c,** Quantification of cell state frequencies from **b**. Approximately 50% of Δ*rpoS* cells were in the heat shock state at LAG10, declining to 10% at LAG30, and nearly undetectable at EXP. **d,** Dot plot showing representative marker genes enriched in the heat shock cluster observed in the Δ*rpoS* strain. **e,** Schematic model summarizing differences in transcriptional trajectories between WT and Δ*rpoS* strains. In WT cells, the transition from stationary phase proceeds through ribosomal biogenesis and respiration. In contrast, Δ*rpoS* cells enter a heat shock state before engaging in ribosome biogenesis. The transient heat shock state may contribute to enhanced persistence in the LAG phase observed in the Δ*rpoS* strain.

To investigate the molecular basis of this phase-specific effect, we profiled the transcriptional cell states of *rpoS*-deficient (Δ*rpoS*) and wild-type (WT) strains at the LAG10 and EXP phases using scRNA-seq. Although Δ*rpoS* cells exhibited markedly reduced antibiotic persistence in STA phase, attempts to profile this condition by scRNA-seq yielded insufficient numbers of mRNA/cell and low cell recovery for inclusion in the final analysis, and we therefore restrict the Δ*rpoS* dataset to LAG10 and EXP phases.

At the LAG10 phase, WT cells initiated ribosome biogenesis, as previously observed. In contrast, approximately 50% of Δ*rpoS* cells exhibited strong upregulation of heat shock genes, including *dnaK*, *groES*, and *groEL* (Fig. 4b–d, Supplementary Fig. 14, Supplementary Tables 8,9). Although *rpoS* is well known for mediating the transition from exponential to stationary phase, it also plays a role during the shift from stationary to lag phase by regulating pathways such as iron uptake^33,34^. Our results suggest that in the absence of *rpoS*, cells may compensate by activating the heat shock response to support the transition from dormancy to active growth. Together with our earlier observation that heat shock activation enhances antibiotic survival, this provides a potential mechanistic explanation for the increased survival of Δ*rpoS* cells in the LAG phase.

To capture the temporal dynamics of this reprogramming by *rpoS* deletion, we also profiled Δ*rpoS* cells 30 minutes post-dilution (LAG30). At this point, their transcriptomes more closely resembled WT LAG10 cells, although ∼10% of cells retained a heat shock signature, indicating delayed or incomplete adaptation (Fig. 4b,c). By the EXP phase, Δ*rpoS* and WT cells showed similar transcriptional profiles dominated by respiration (Fig. 4b,c), consistent with their comparable levels of antibiotic persistence in this phase.

Altogether, these findings demonstrate that perturbation of *rpoS* can alter pre-treatment cell states, a potential mechanism through which the level of antibiotic persistence is modulated. While RpoS is classically associated with stress protection and has been implicated in persistence, our results reveal that its effects are strongly context dependent. In the LAG phase, RpoS facilitates the transition from starvation adaptation to ribosome biogenesis; its absence instead triggers a heat shock response and paradoxically increases persistence (Fig. 4e). These results provide a potential mechanistic basis for the growth phase-specific effects of RpoS on persistence and underline the limitations of gene-targeted approaches for modulating antibiotic persistence.

### Nutrient shift reprograms stationary phase cells and restores antibiotic sensitivity

Having shown that genetic perturbation of pre-treatment cell states alters antibiotic survival, we hypothesized that non-genetic interventions, such as nutrient shifts, could similarly modulate transcriptional states and sensitize tolerant cells. This concept has been proposed by prior work showing that certain carbon sources can activate aminoglycoside uptake and killing of *E. coli* persisters via restoration of proton motive force^35^. However, these effects were largely limited to aminoglycosides and varied depending on bacterial species and nutrient type. Moreover, past efforts focused on bulk phenotypes and did not investigate how nutrient supplementation alters the transcriptional landscape or interacts with antibiotics beyond aminoglycosides. Here, we tested whether specific nutrient inputs could shift bacterial cell states and alter antibiotic survival under meropenem and ciprofloxacin. If successful, this strategy could provide a generalizable way to enhance antibiotic efficacy by shifting bacterial populations into more vulnerable physiological states.

To evaluate this, we focused on STA cells, which exhibited the highest levels of antibiotic tolerance. We screened a variety of nutrients, including carbon sources, nitrogen sources, amino acids, and metals, by supplementing them individually into spent medium (filtered from overnight MHB cultures), and assessed survival under meropenem and ciprofloxacin treatment (Supplementary Fig. 15). For meropenem-treated STA cells, glucose supplementation reduced survival by roughly 1000 folds after 4-hour treatment (Fig. 5a). For ciprofloxacin-treated STA cells, ferrous sulfate (FeSO_4_) reduced survival by approximately 30–100 folds after 4-hour treatment (Fig. 5b). Notably, glucose had no effect under ciprofloxacin, and FeSO_4_ had no effect under meropenem (Supplementary Fig. 15), suggesting that these two interventions sensitize cells through distinct mechanisms. Importantly, we confirmed that glucose or FeSO_4_ alone did not affect survival in the absence of antibiotics (Supplementary Fig. 16), indicating that their effects depend on the antibiotic used.

**Figure 5.**
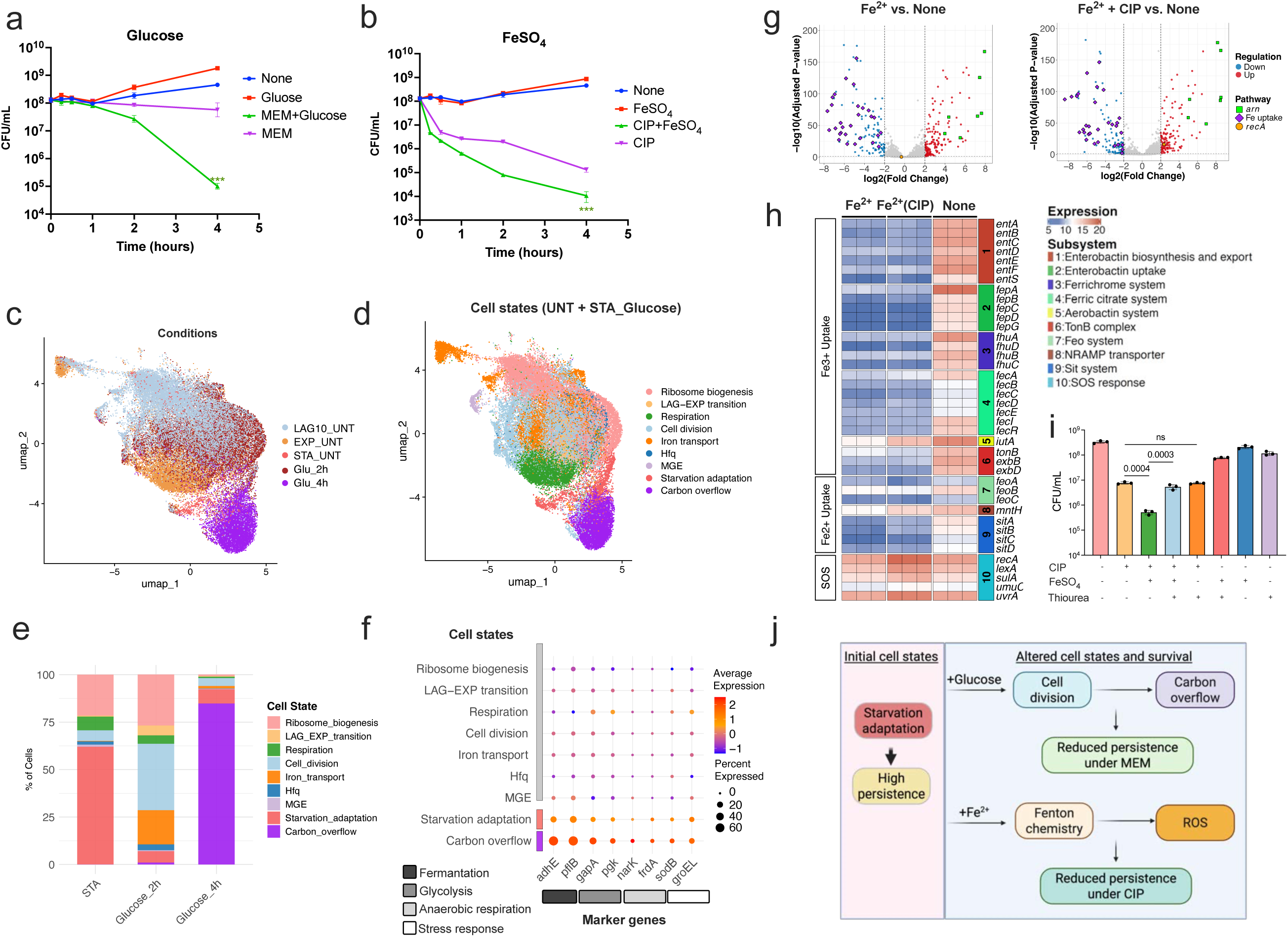
Nutrient shift reprograms stationary phase cells and restores antibiotic sensitivity. **a,** Killing dynamics of stationary-phase (STA) *K. pneumoniae* MGH66 cells transferred to spent medium with or without 20 mM glucose, and treated with meropenem (MEM) (2 µg/ml) as indicated. Glucose supplementation significantly enhanced meropenem-mediated killing compared to meropenem alone (n = 3, *p* = 0.0007, Student’s *t*-test, 4 hours). **b,** Killing dynamics of STA cells transferred to spent medium with or without 10 mM FeSO_4_, and treated with ciprofloxacin (CIP) (2.5 µg/ml) as indicated. FeSO_4_ supplementation significantly enhanced ciprofloxacin-mediated killing compared to ciprofloxacin alone (n = 3, *p* = 0.0014, Student’s *t*-test, 4 hours). Experiments in **a** and **b** were performed with three biological replicates. Curves represent the mean of three replicates; error bars denote standard deviation. **c,** UMAP embedding of single-cell transcriptomes from five conditions: three untreated (UNT) samples at early lag (10 minutes post-transfer to fresh medium, LAG10_UNT), exponential (EXP_UNT), and stationary phases (STA_UNT), and two glucose-treated STA samples (STA_glucose) collected at 2 hours (Glu_2h) and 4 hours (Glu_4h) post-supplementation. Each condition includes two biological replicates (n = 2). **d,** Same UMAP as in **c**, colored by transcriptional cell states. In addition to the eight shared states identified in untreated cells, glucose-treated STA cells entered a carbon overflow state at 4 hours post glucose-supplementation. **e,** Bar plot showing the distribution of cell states across three conditions: STA, glucose-treated STA samples collected at 2 hours (Glucose_2h) and 4 hours (Glucose_4h) post-supplementation. Percentages represent the average from two biological replicates (n = 2). **f,** Dot plot showing representative marker genes for the carbon overflow state identified in glucose-supplemented STA cells. **g,** Volcano plots of bulk RNA-seq from STA cells supplemented with Fe²^+^ (10 mM), with or without ciprofloxacin (2.5 µg/ml) treatment. Fe²^+^ alone upregulated the *arn* operon and downregulated iron uptake genes. Additional ciprofloxacin treatment induced *recA*, while maintaining iron regulation changes. Genes with log_2_ fold change > 2 and adjusted *p* < 0.05 are highlighted. Three biological replicates were included. **h,** Heatmap showing gene-level expression in Fe²^+^- and Fe²^+^ + ciprofloxacin-treated cells. Nearly all iron uptake pathways were downregulated following Fe²^+^ supplementation, while ciprofloxacin additionally induced the SOS response. **i,** Supplementation of thiourea (200 mM) significantly attenuated the Fe²^+^-enhanced (10 mM FeSO_4_) killing effect of ciprofloxacin (2.5 µg/ml), suggesting a role for reactive oxygen species (ROS). Three biological replicates (n = 3) were performed. CFU was quantified at 4 hours post ciprofloxacin treatment and Student’s *t*-test was performed for the statistical analysis. **j,** Schematic model summarizing the distinct mechanisms by which glucose and Fe²^+^ supplementation restore antibiotic sensitivity in STA cells. Glucose shifts cells from stationary phase into active division (2 hours) and carbon overflow metabolic states (4 hours), enhancing meropenem killing by reactivating the expression of drug targets. Fe²^+^ triggers Fenton chemistry, converting Fe²^+^ to Fe³^+^ and generating ROS that enhance ciprofloxacin-mediated killing. In both cases, altered initial cell states interact with drug-specific mechanisms to modulate antibiotic susceptibility.

To uncover how glucose sensitizes stationary-phase cells to meropenem, we performed scRNA-seq on STA cells transferred into spent medium supplemented with 20 mM glucose and sampled them at 2 and 4 hours post-transfer. By 2 hours, cells had re-entered growth, exhibiting active DNA replication and cell wall synthesis (Fig. 5c–e; Supplementary Fig. 17; Supplementary Tables 10,11). By 4 hours, ∼80% of cells transitioned into a transcriptional state of carbon overflow^36,37^, characterized by mixed metabolic reprogramming and stress responses (Fig. 5c–e), including upregulation of fermentation genes, glycolysis genes, anaerobic respiration genes, and stress response pathways (Fig. 5f, Supplementary Tables 10,11). Because replicates differed substantially in recovered cell numbers for this condition, we restrict interpretation to the predominant transcriptional programs rather than quantitative estimates of rare states. These findings indicate that glucose supplementation drives a growth trajectory distinct from that induced by fresh medium, which primarily activates aerobic respiration (Fig. 1g). Instead, under carbon excess but limitation in other nutrients, cells engage alternative metabolic programs to support growth, rendering them more susceptible to meropenem-mediated killing.

We next investigated how FeSO_4_ enhances ciprofloxacin killing. Unlike glucose supplementation, which allowed us to profile transcriptional states using scRNA-seq, FeSO_4_-treated STA cells exhibited low RNA content, and we were unable to recover enough high-quality cells for single-cell analysis. We therefore turned to bulk RNA-seq, which offers greater sensitivity. In the absence of antibiotic, FeSO_4_ supplementation led to robust upregulation of the *arn* operon, a pathway involved in lipid A modification and known to respond to Fe²^+^. We also observed significant downregulation of ferric (Fe³^+^) iron uptake systems, suggesting a shift in iron homeostasis (Fig. 5g,h; Supplementary Tables 12,13). Under ciprofloxacin treatment, these transcriptional changes persisted, along with a marked induction of the SOS response (Fig. 5g,h). The downregulation of Fe³^+^ uptake pathways suggests intracellular Fe³^+^ accumulation, potentially resulting from Fe²^+^ oxidation following supplementation. We thus hypothesized that the conversion of Fe²^+^ to Fe³^+^ via Fenton chemistry generates reactive oxygen species (ROS), thereby potentiating antibiotic lethality^38^. Supporting this model, the addition of thiourea, a ROS scavenger, attenuated the killing effect (Fig. 5i). Notably, thiourea alone did not affect ciprofloxacin-mediated killing, suggesting that in the absence of Fe²^+^, ciprofloxacin acts through a ROS-independent mechanism in STA cells. Together, these results reveal a mechanistic basis for how FeSO_4_ potentiates ciprofloxacin activity against antibiotic-tolerant STA cells through Fe^2+^-driven oxidative stress that synergizes with drug-induced damage.

Altogether, these results demonstrate that nutrient shifts can serve as an effective strategy to sensitize antibiotic-tolerant cells. Notably, the mechanism of sensitization depends on both the antibiotic used and how nutrient-induced transcriptional changes intersect with the drug’s mode of action (Fig. 5j).

## Discussion

By applying bacterial scRNA-seq, we were able to directly profile the heterogeneous responses that linked to antibiotic survival, providing insight into early programs that may precede persister formation. This approach revealed that pre-treatment cell states play a critical role in shaping transcriptional reprogramming profiles, which in turn determine survival outcomes under antibiotic stress. Using growth phase as a biologically meaningful axis of heterogeneity, we found that *K. pneumoniae* cells can engage multiple, distinct survival pathways simultaneously. These divergent reprogramming pathways are largely dictated by the interplay between a cell’s pre-treatment transcriptional state and the antibiotic’s mechanism of action. Genetic and environmental perturbations, such as *rpoS* deletion and nutrient supplementation, shifted pre-treatment cell states and thereby redirected reprogramming and altered persistence frequencies. Notably, the effect of *rpoS* deletion was conditional on growth phase, illustrating how redundancy and plasticity within stress responses buffer against single-gene perturbations. Together, these results provide direct mechanistic evidence linking pre-treatment cell states to antibiotic-induced transcriptional reprogramming and persistence, framing persistence as a systems-level property shaped by heterogeneity and redundancy.

### Profiling bacterial heterogeneity and its biological significance

Our analysis revealed that *K. pneumoniae* cells exposed to antibiotics did not respond uniformly but instead followed multiple transcriptional responses, each associated with distinct survival outcomes. For example, under meropenem treatment, STA-phase cells reprogrammed minimally and survived; LAG-phase cells activated heat shock; and EXP-phase cells diversified into subpopulations that engaged upregulation of *osmB*, starvation adaptation, or heat shock responses. These divergent responses were not random but constrained by baseline transcriptional states. Importantly, multiple responses could support cell survival within the same isogenic population at the same growth phase (Fig. 2g,h), indicating that survival is not determined by a single conserved program. Moreover, deletion of *rpoS* had conditional effects on persistence (Fig. 4a), illustrating that no single gene is determinative and that redundancy within stress responses contributes to survival. Together, these findings demonstrate the biological significance of transcriptional heterogeneity shaped by pre-treatment states, explaining why persistence remains robust across conditions and resistant to single-gene perturbations.

A limitation of the current study is reduced sampling depth for stationary-phase cells, particularly under antibiotic treatment, reflecting the low RNA content of these cells and current technical constraints of bacterial scRNA-seq. While additional analyses with relaxed filtering thresholds recovered consistent transcriptional states, rare subpopulations may remain undetected. Similarly, stationary-phase cells under the FeSO_4_-treated conditions and in the Δ*rpoS* strain exhibited markedly reduced numbers of detected genes per cell despite comparable sequencing depth, and we were unable to recover sufficient cells for downstream analysis. These observations are consistent with biological states characterized by low recoverable mRNA, although technical sensitivity remains a general challenge for bacterial scRNA-seq. Further advances in sensitivity will be required to resolve such low-transcription states.

While growth phase served as a practical axis of heterogeneity in this study, we also observed substantial transcriptional variation within each phase. The functional relevance of this within-phase heterogeneity remains unresolved. Addressing it will require lineage tracing and advanced computational modeling that directly link pre-treatment cell state to cell fate^39^, approaches that are not yet available for bacterial scRNA-seq. Although we defined transcriptional states as discrete clusters, these likely lie along a continuum of regulatory and physiological transitions. Our scRNA-seq analysis offers snapshots of this landscape, but a fuller understanding of persistence will require time-resolved approaches that can capture continuous trajectories and dynamic reprogramming at single-cell resolution.

### Modulating cell states to overcome persistence

Our findings demonstrate that rationally shifting bacterial cell states can reprogram transcriptional states and reshape antibiotic susceptibility. While previous studies have proposed targeting specific genes or pathways to overcome persistence, our results caution against such gene-centric approaches. For example, deletion of *rpoS*, a conserved global stress response regulator, produced divergent effects on persistence depending on growth phase (Fig. 4a). This context-dependent behavior underscores the limitations of single-gene interventions and supports the need for a systems-level strategy to modulate bacterial cell states.

Rather than focusing on individual persistence factors, we propose that interventions should aim to reshape the overall transcriptional landscape to drive bacterial populations into vulnerable states. Environmental perturbations, such as nutrient supplementation, may offer one such route. While further work is needed to translate this strategy into clinical settings, cell-state-based modulation may offer a broadly effective route to overcome antibiotic persistence.

### Toward a broader and *in vivo* understanding of antibiotic persistence

Our findings, while derived from a clinical isolate of *K. pneumoniae* from a urinary tract infection, a context frequently associated with recurrence and persistence, likely reflect conserved principles that extend beyond this strain and species. The transcriptional states we identified, both before and after antibiotic exposure, involve core metabolic pathways and conserved stress responses such as the RpoS-mediated general stress response and the SOS response. These are well-characterized pathways with established roles in persistence across multiple species. For instance, the conditional effect of *rpoS* deletion on persistence was not only observed in *K. pneumoniae* (Fig. 4a) but also validated in *E. coli* (Supplementary Fig. 13), and is consistent with previous reports in *E. coli* and *Pseudomonas*^16^, suggesting this phenomenon is not strain-specific. Together, these findings suggest that our observations are not unique to a particular isolate but instead represent broadly relevant cell-state features with implications for persistence across species.

Our experiments were conducted under well-controlled *in vitro* conditions, enabling precise dissection of how pre-treatment transcriptional states shape antibiotic responses. In contrast, *in vivo* environments introduce additional complexity, including spatial organization, immune pressure, and polymicrobial interactions, that may reshape bacterial physiology and cell-state dynamics. Therefore, interpreting bacterial single-cell responses in such complex environments requires robust reference frameworks.

Moving forward, the cell-state markers defined here provide a foundational resource for annotating bacterial states *in vivo* and for generating mechanistic hypotheses about persistence. Together, this work offers both mechanistic insight and a technical roadmap for future studies across diverse pathogens and infection contexts, advancing efforts to rationally reprogram bacterial cell states and improve antibiotic efficacy.

## Methods

### Bacterial strains and growth conditions

All scRNA-seq experiments are listed in Supplementary Table 14. The bacterial strains used in this study are listed in Supplementary Table 15. *K. pneumoniae* MGH66^40^ served as the parental strain for generating derivatives. *K. pneumoniae* strains were routinely grown in Mueller-Hinton Broth (MHB) at 37°C with shaking at 220 rpm. Competent cells of *E. coli* NEB 10-beta, employed for plasmid cloning and propagation, were purchased from New England Biolabs (NEB). *E. coli* K-12 wild-type and its derived Δ*rpoS* mutant strains were obtained from the Keio collection^41^.

To investigate transcriptional heterogeneity across bacterial growth phases and with antibiotic stress, *K. pneumoniae* cells were collected at distinct growth stages for scRNA-seq. To obtain lag-phase cells, stationary-phase cultures were diluted 1:100 into 30-ml fresh MHB and grown at 37°C for 10 min before harvesting. For exponential-phase cells, stationary-phase cultures were first diluted 1:100 into fresh MHB and grown to an OD_600_ of 0.2. The cultures were then subjected to two additional rounds of 1:100 back-dilution into 30-ml fresh MHB, each time grown to an OD_600_ of 0.2. Cells were collected after the final dilution when the OD_600_ reached 0.2. Stationary phase cells were obtained by inoculating glycerol stocks into fresh MHB and incubating for approximately 16 hours. The resulting populations were designated as lag-phase (LAG10), exponential-phase (EXP), and stationary-phase (STA) cells, respectively. To characterize the transcriptional profiles of nutrient-shifted STA cells, STA cultures were diluted 1:100 into nutrient-spent medium supplemented with 20 mM glucose and grown for 2 or 4 hours. Nutrient-spent medium was obtained from the supernatant of overnight cultures and sterilized using a 0.22 µm vacuum driven filtration system (Millipore, S2GPU02RE). For the antibiotic-treated samples, cells from each growth phase were incubated with 2 μg/ml meropenem (Sigma-Aldrich, 1392454) or 2.5 μg/ml ciprofloxacin (Sigma-Aldrich, 17850) at 37°C with shaking for the indicated time periods. After antibiotic exposure, cells were collected by centrifugation at 4°C for 15 min at 4,500g and immediately fixed using 4% formaldehyde in PBS, incubate overnight at 4°C, then processed for subsequent scRNA-seq experiments.

### Cell fixation and permeabilization

The scRNA-seq protocol used in this study (BacDrop_V2) was optimized by replacing the second-round barcoding strategy from BacDrop^23^ with the thermo-ligation based in-droplet barcoding, as validated and described by previous studies^24,42^. Details of the workflow are shown in Supplementary Fig. 1. Key reagents or kits used in this study are listed in Supplementary Table 16.

The cell fixation and permeabilization were performed following our previously established protocol. Briefly, Bacterial cells (∼10⁹ to10¹⁰) grown under specified conditions were collected by centrifugation at 4,500g for 15 min at 4°C and resuspended in 7 ml ice-cold fresh 4% formaldehyde in 1× PBS. Fixation was carried out overnight at 4°C with gentle shaking. Following fixation, cells were pelleted by centrifugation at 4,500g for 15 min at 4°C and resuspended in 700 μl PBS-RI (1× PBS supplemented with 0.1 U/ml NxGen RNase Inhibitor) in a nuclease-free 1.5 ml microcentrifuge tube (Axygen, 14324061). All subsequent centrifugations were performed at 12,700 rpm for 4 min at 4°C. Cells were pelleted and resuspended in 700 μl PBS-RI, centrifuged again under the same conditions, and resuspended in 350 μl PBS-RI. Then, 350 μl of 100% ethanol was added to the cell suspension and mixed gently by pipetting. Immediately afterward, cells were washed twice with 700 μl PBS-RI. After the second wash, cells were resuspended in 1 ml of ice-cold 100 mM Tris-HCl-RI (100 mM Tris-HCl pH 7.5 supplemented with 100 U/ml SUPERase-In RNase Inhibitor). Cells were diluted 1:100 in PBS and counted using a hemocytometer (Bulldog-Bio, DHC-N420) to determine its concentration.

For each permeabilization reaction, up to 1 × 10⁸ cells were resuspended in 250 μl of 0.04% Tween-20 in PBS-RI and incubated on ice for 3 min. Subsequently, 1 ml of ice-cold PBS-RI was added, and cells were pelleted by centrifugation. The pellet was resuspended in 400 μl of lysozyme solution (100 mM Tris-HCl pH 8.0, 50 mM EDTA, 250 U/ml SUPERase-In RNase Inhibitor, 2.5 mg/ml lysozyme) and incubated at 37°C for 15 min. Following incubation, 1 ml of PBS-RI was added, cells were pelleted by centrifugation and resuspended in 175 μl of PBS-RI. Cell concentration was determined using a hemocytometer.

### In situ rRNA depletion and gDNA digestion

Immediately following cell permeabilization, 4 × 10^7^ cells were pelleted and resuspended in 11 μl of nuclease-free water. The rRNA depletion was carried out using a thermostable RNase H-based protocol with the NEBNext Bacterial rRNA Depletion Kit as described previously^23^. Specifically, 2 μl of NEBNext rRNA Depletion Solution and 2 μl of Probe Hybridization Buffer were added to the cells on ice. Hybridization was performed using the PCR cycler with following thermal profile (lid set to 55°C): 50°C for 2 min, then ramped down to 22°C at 0.1°C/sec, followed by a 5-min hold at 22°C. Immediately after hybridization, RNase H digestion was initiated by adding 2 μl of RNase H Reaction Buffer, 2 μl of Thermostable RNase H, and 1 μl of nuclease-free water. The reaction was incubated at 50°C for 30 min (lid at 55°C).

Following rRNA depletion, cells were centrifuged and resuspended in 10 μl of DNase-RI buffer (1 μl 10× DNase I Reaction Buffer, 1 μl DNase I, 0.5 μl SUPERase-In RNase Inhibitor, and 7.5 μl nuclease-free water). The mixture was incubated at room temperature for 30 min. To terminate DNase activity, 1 μl of Stop Solution was added, followed by incubation at 50°C for 10 min. Afterward, cells were centrifuged and washed twice with 100 μl of PBS-RI. The final cell pellet was resuspended in 20 μl of 0.5× PBS-RI and used immediately for in-cell reverse transcription.

### Modifications to the previously published BacDrop protocol^23^

To improve barcoding efficiency relative to the previously published BacDrop_V1 workflow, we implemented a modified barcoding scheme in BacDrop_V2 that changes how round-2 barcodes (CB2) are appended. In BacDrop_V1, in-cell reverse transcription (RT) is performed using RT primers carrying the round-1 barcode (CB1), followed by in-cell 3′ poly(A) tailing of the newly synthesized cDNA. Cells are then encapsulated into droplets, where oligo(dT)-primed second-strand synthesis generates a cDNA duplex whose 5′ end can anneal to the 10x gel-bead oligonucleotide carrying a unique barcode; polymerase extension subsequently copies the bead barcode onto the molecule to introduce the round-2 barcode (CB2). In BacDrop_V2, we retained in-cell RT and CB1 incorporation but redesigned the RT primer architecture (Supplementary Table 17) such that the cDNA produced in cells is compatible with a ligation-based in-droplet cell barcoding (CB2). Following in-cell RT, we omitted the in-cell poly(A) tailing step in BacDrop_V1 and proceeded directly to droplet encapsulation using the same ATAC-seq kit from 10x Genomics. Within droplets, CB2 was introduced by a bridge-oligo-mediated (Supplementary Table 18) ligation reaction that directly joins the cDNA (via the RT-primer encoded 5′ end modified with a phosphate group) to the 10x ATAC-seq gel-bead oligonucleotide containing the bead-specific barcode, thereby replacing the V1 polymerase-driven CB2 barcoding step based on second-strand synthesis. This ligation-based round-2 barcoding strategy and primer/bridge-oligo design principles were adapted from the published SciFi-RNA-seq and M3-seq protocols^24,42^. After droplet barcoding, cDNA from all droplets was pooled and purified, poly(A) tailing was performed at the pooled stage, and oligo(dT)-primed second-strand synthesis was carried out prior to downstream library construction. Subsequent steps followed the BacDrop V1 workflow, including tagmentation to add Illumina adapters and PCR amplification for sequencing-ready libraries. We validated the modified workflow by comparing pseudo-bulk profiles derived from BacDrop V2 to matched bulk RNA-seq data and observed high concordance, indicating that the updated barcoding strategy did not introduce appreciable bias (Supplementary Fig. 1b,c). Detailed protocols and V2-specific modifications are provided below.

### In-cell reverse transcription and round 1 barcoding

The first round of cell barcoding and multiplexing was conducted via RT in 96-well plates. Compared to previously described BacDrop protocol, we modified our RT primers to optimize the round 1 and round 2 barcoding efficiency. The modified RT primers (Supplementary Table 17) containing both a unique molecular identifier (UMI) and a round 1 cell barcode (CB1) were custom-synthesized (Integrated DNA Technologies) at 100 μM and diluted to 5 μM in a 96-well PCR plate (Axygen, PCR-96-HS-C). Each well contained a RT primer containing distinct, well-specific CB1 sequences, ensuring that every well corresponded to a unique CB1 barcode. A phosphate group was added the 5’ of each RT primer for the ligation step in round 2 barcoding. Each well of the RT plate was preloaded with 5 μl of a distinct primer (5 μM). Cells previously depleted of rRNA and genomic DNA were diluted in 0.5× PBS-RI and added at 2 μl per well to the primer-containing plates. The cell-primer mix was incubated at 55°C for 5 min to facilitate primer annealing, followed by immediate cooling on ice. An RT master mix consisting of 0.5 μl DTT (100 mM), 0.5 μl dNTP mix (10 mM each), 0.5 μl SUPERase-In RNase inhibitor, 2 μl Maxima RT buffer, and 0.5 μl Maxima H Minus reverse transcriptase was added to each well. Reverse transcription was carried out in the PCR cycler under the following program (lid at 60°C): Initial incubation at 22°C for 30 min and 50°C for 10 min, followed by three thermocycling rounds of 8°C for 12 s, 15°C for 45 s, 20°C for 45 s, 30°C for 30 s, 42°C for 2 min, 50°C for 3 min. The reaction ended with a final 5-min incubation at 50°C and holding at 4°C. Following RT, cells were pelleted and washed 2 times with PBS-RI, then resuspended in 20 μl nuclease-free water supplemented with 200 U/ml SUPERase-In RNase inhibitor (H_2_O-RI). Cell density was subsequently determined using a hemocytometer.

### Droplet generation

The round 2 barcoding was modified from the previous described BacDrop_V1 protocol to increase the coverage and barcoding efficiency. Specifically, we replaced the previous second strand synthesis protocol with a thermo-ligation protocol as described in several other established scRNA-seq protocols. For droplet formation, the Chromium Next GEM Chip H and the Single Cell ATAC Library & Gel Bead Kit were utilized following the instructions with minor adjustments. Unused wells on the chip were filled with 50% glycerol, specifically, 70 μl in row 1, 50 μl in row 2, and 40 μl in row 3. Cells prepared for encapsulation were diluted with H_2_O-RI to a final volume of 63 μl. Just before loading onto the chip, the cell suspension was mixed with a ligation master mix consisting of 11.5 μl Ampligase buffer, 2.3 μl 100 μM bridge oligo (BOa, Supplementary Table 18), 2.3 μl Ampligase, and 1.5 μl Reducing Agent B (Single Cell ATAC Library & Gel Bead Kit). A total of 70 μl of the resulting mixture was loaded into row 1 of the chip. 50 μl of gel beads embedded with round 2 barcodes (CB2) and 40 μl of partitioning oil were loaded into rows 2 and 3 of the Chromium chip, respectively. The chip was then processed using the Chromium iX controller, generating an emulsion of approximately 100 μl containing encapsulated single cells.

### Round 2 barcode ligation

Following droplet generation, the emulsion was evenly distributed into four individual PCR tubes (approximately 25 μl per tube) using TempAssure 8-tube PCR strips (USA Scientific, 14024780). To each tube, 25 μl of FC-40 oil was first added to the bottom, followed by 50 μl of mineral oil layered on top of the emulsion, forming a biphasic mixture. Tubes were then subjected to ligation in a thermal cycler using the following program (lid at 105°C): 12 cycles of 95°C for 30 s and 59°C for 2 min, followed by hold at 15°C. After ligation, samples were left at room temperature for 15 min for better phase separation. The upper mineral oil and lower FC-40 oil layers were carefully removed by pipetting, ensuring the droplet-containing middle layer remained undisturbed.

### Breaking emulsions and cDNA purification

All reagents required for breaking emulsions were sourced from the Single Cell ATAC Library & Gel Bead Kit. To disrupt the emulsion in each PCR tube, 32 μl of Recovery Agent was added at room temperature. Tubes were gently inverted ten times and briefly spun down to ensure complete phase separation. The lower pink Recovery Agent and partitioning oil layer was carefully removed. cDNA purification was initiated using Dynabeads MyOne SILANE. Specifically, for each sample, 45.5 μl Cleanup Buffer, 2 μl Dynabeads, 1.25 μl Reducing Agent B, and 1.25 μl nuclease-free water were added. After a 10-min incubation at room temperature with gentle mixing, tubes were placed on a magnetic rack, and beads were washed twice using freshly prepared 80% ethanol. After the second ethanol wash was removed, cDNA was eluted in 40 μl of a custom elution buffer (98 μl Buffer EB, 1 μl 10% Tween-20, and 1 μl Reducing Agent B). The eluate was transferred to a new PCR tube for further purification using 1.25× SPRIselect beads, and the final cDNA was recovered in 43 μl nuclease-free water.

### TdT tailing and second strand of cDNA synthesis

Compared to the original BacDrop_V1 protocol, the terminal deoxynucleotidyl transferase (TdT) tailing and second-strand synthesis steps were performed after cDNA purification from the second-round droplet barcoding. In the modified workflow, TdT tailing was performed on purified cDNA rather than on permeabilized cells. A total of 43 μl of barcoded cDNA (containing CB1 and CB2) was mixed with 5 μl of 10× TdT buffer, 1 μl of 100 mM dATP, and 1 μl of TdT enzyme (20 U/μl), resulting in a final reaction volume of 50 μl. The reaction was incubated at 37°C for 30 min. To terminate the reaction, 10 μl of 0.2 M EDTA (pH 8.0) was added, followed by a 10-min incubation at room temperature. The reaction mixture was then purified using a 2× SPRIselect bead cleanup and eluted in 44 μl of nuclease-free water.

Second-strand cDNA synthesis was carried out using the KAPA HiFi HotStart ReadyMix. Each 100 μl reaction included 44 μl of purified single-stranded cDNA, 50 μl of 2× KAPA HiFi HotStart ReadyMix, 3 μl of 10 μM SMRT_dT primer, and 3 μl of 10 μM P5 primer. The PCR cycling protocol was as follows (lid at 105°C): 95°C for 30 s; 39°C for 5 min; four cycles of 65°C for 10 min, 98°C for 20 s, 62°C for 15 s, and 72°C for 2 min; followed by 72°C for 5 min and a final hold at 10°C. The amplified product was then purified using a 2× SPRIselect bead cleanup and eluted in 50 μl of nuclease-free water. To assess cDNA concentration, 2 μl of each sample was measured using a Qubit fluorometer with the Qubit 1× dsDNA High Sensitivity (HS) Assay Kit. Samples yielding decent DNA concentration were selected for downstream library construction.

### Enrichment of cDNA

Prior to cDNA amplification, a quantitative PCR (qPCR) assay was performed to determine the optimal number of enrichment cycles. The reaction consisted of 1 μl cDNA template, 5 μl KAPA HiFi HotStart ReadyMix, 0.3 μl each of P5 (10 μM) and SMRT_PCR primers (10 μM) (Supplementary Table 18), 2 μl of 5× SYBR Green dye, and 1.4 μl nuclease-free water. qPCR was run on a real-time thermal cycler (CFX Duet, BioRad) using the following cycling conditions: initial denaturation at 98°C for 2 min; 30 cycles of 98°C for 20 s, 67°C for 20 s, and 72°C for 3 min; followed by a final extension at 72°C for 5 min and hold at 4°C. The cycle threshold (Ct) corresponding to the early exponential phase of amplification was used to guide subsequent cDNA enrichment. For large-scale amplification, 44 μl of cDNA was combined with 50 μl of 2× KAPA HiFi HotStart ReadyMix, 3 μl each of P5 and SMRT_PCR primers (10 μM) to a total volume of 100 μl. The PCR was performed using the same thermal cycling program as in the qPCR, applying the Ct-derived cycle number. Following amplification, the product was cleaned up using a 0.6× SPRIselect bead purification protocol and eluted in 30 μl of nuclease-free water for subsequent sequencing library preparation. cDNA concentration was quantified using a Qubit fluorometer in combination with the Qubit 1× dsDNA High Sensitivity (HS) Assay Kit.

### Illumina library preparation for sequencing

Sequencing libraries were prepared using the Illumina Nextera XT DNA Library Prep Kit with protocol adjustments. For each 20 μl tagmentation reaction, 1 to 2 ng of enriched cDNA was mixed with 5 μl of Amplicon Tagment Mix (ATM) and 10 μl of Tagment DNA Buffer (TD). The mixture was incubated at 55°C for 5 min in a thermocycler, followed by a hold at 10°C. Once the reaction reached 10°C, 5 μl of Neutralize Tagment Buffer (NT) was added immediately. The solution was mixed by pipetting up and down 10 times and incubated at room temperature for 5 min before proceeding to PCR amplification. Next, 5 μl of P5 primer (5 μM) (Supplementary Table 18), 5 μl of Index 1 primer (N7**) (FC-131-2001, Illumina), and 15 μl of Nextera PCR Master Mix (NPM) were added to the 25 μl tagmented mixture to form a 50 μl PCR reaction. The thermal cycling program included an initial extension at 72°C for 3 min and denaturation at 95°C for 30 s, followed by 5 cycles of 95°C for 10 s, 55°C for 30 s, and 72°C for 30 s, and a final extension at 72°C for 5 min. The PCR tubes were immediately placed on ice to avoid over-amplification.

To determine the optimal number of additional amplification cycles, a qPCR assay was set up using 5 μl of the partially amplified library, 3 μl of NPM, 1 μl each of P5 and Index 1 primers (5 μM), 3 μl of 5× SYBR Green, and 2 μl of nuclease-free water. The qPCR was performed under the same cycling conditions. The cycle number corresponding to the midpoint of the exponential amplification curve was selected for the remaining library amplification. The remaining 45 μl of the initial PCR mixture was subjected to additional amplification using the cycle count determined by qPCR. Final libraries were purified with a 1× SPRIselect bead cleanup and eluted in 25 μl of nuclease-free water. Library concentrations were quantified using the Qubit 1× dsDNA High Sensitivity (HS) Assay Kit and a Qubit fluorometer.

### Illumina sequencing of the BacDrop_V2 library

Prepared libraries were diluted to the appropriate concentrations and sequenced on the Illumina NovaSeq X plus platform using standard Illumina sequencing primers. Sequencing was performed with the following read configuration: Read 1: 60 bp, Read 2: 40 bp, Index 1: 8 bp, and Index 2: 30 bp.

### Antibiotic killing curve assay

To investigate antibiotic persistence across growth phases, bacterial cultures were grown in MHB to stationary, lag, or exponential phase as described above. A lethal final concentration of either meropenem (2 μg/ml) or ciprofloxacin (2.5 μg/ml) was added to the culture and incubate at 37°C with shaking. At the specified time points post-treatment, 1 ml of culture was harvested, centrifuged (12,700 rpm, 3 min), resuspended in 100 μl PBS, and subsequently serially diluted in PBS. Ten microliters of each were spotted onto LB agar plates. After incubation at 37°C for 16-18 hours, colony-forming units (CFUs) were counted to quantify survived bacteria. Survival rates were calculated by normalizing CFUs at each time point to the initial CFUs at time zero (before antibiotic exposure). Each killing assay was performed with at least three biological replicates.

### Construction of GFP reporter strains

GFP reporter plasmids were constructed by inserting the target promoter into the pUA139^43^ vector upstream of the *gfp* reporter gene. Briefly, the plasmid backbone was amplified by PCR using Q5 High-Fidelity 2× Master Mix with primers PMSJ060 and PMSJ061 (Supplementary Table 18). Synthesized promoter of *lpp1*, *osmB*, *rpoS*, *recA*, *dnaK*, and *fusA* genes (Supplementary Table 18), each containing homologous overlaps with the plasmid backbone, were assembled with the amplified pUC139 fragment using the NEBuilder HiFi DNA Assembly Cloning Kit. The assembled constructs were transformed into NEB 10-beta for propagation. Plasmid integrity and sequence correctness were verified via whole plasmid sequencing. The validated reporter plasmids (Supplementary Table 15), pUA139_P*lpp1_gfp*, pUA139_P*osmB_gfp*, pUA139_P*rpoS_gfp*, pUA139_P*recA_gfp*, and pUA139_P*dnaK_gfp*, were subsequently transformed into *K. pneumoniae* MGH66 WT by electroporation as previously described. This yielded the reporter strains (MGH66:pUA139_P*lpp1_gfp*, MGH66:pUA139_P*osmB_gfp*, MGH66:pUA139_P*rpoS_gfp*, MGH66:pUA139_P*recA_gfp*, and MGH66:pUA139_P*dnaK_gfp*) for flow cytometric sorting and analysis.

### Flow cytometry analysis and cell sorting for antibiotic killing assay

Exponential-phase cultures of reporter strains were prepared as described above in MHB medium supplied with 50 μg/ml kanamycin (Sigma-Aldrich, K1377) to maintain the reporter plasmid. Upon reaching exponential phase, the reporter strains *K. pneumoniae* MGH66: P*osmB:gfp*, MGH66: P*rpoS:gfp*, and MGH66: P*fusA:gfp* were treated with 2 μg/ml meropenem for 10 min. Following antibiotic exposure, cells were diluted in PBS staining buffer (PBS containing DAPI) to a final concentration of 10^6^ cells/ml, and immediately analyzed by flow cytometry. For persister assay under the treatment of meropenem, GFP-low and GFP-high subpopulations were isolated by fluorescence-activated cell sorting (FACS). From each sorted population, approximately 10^6^ cells were collected into 2 ml MHB medium or MHB medium supplemented with 2 μg/ml meropenem. From the antibiotic-free MHB medium, a 50 μl aliquot of sorted cells was immediately serially diluted and plated on LB agar to determine the initial CFUs. The cells sorted into MHB medium containing meropenem were incubated at 37°C with shaking for 4 hours, washed, serially diluted in PBS, and plated on LB agar. CFUs were enumerated to quantify surviving bacteria. Three biological replicates were performed in each experiment.

For flow cytometry analysis of stationary-phase cells exposed to antibiotics, *K. pneumoniae* MGH66 carrying the reporter plasmids pUA139_P*lpp1_gfp* or pUA139_P*recA_gfp* was cultured to stationary phase in MHB supplemented with kanamycin (50 μg/ml), as described above. Stationary-phase MGH66:pUA139_P*lpp1_gfp* and MGH66:pUA139_P*recA_gfp* cells were exposed to meropenem (2 μg/ml) or ciprofloxacin (2.5 μg/ml), respectively, for 30 min. Following treatment, cells were washed, resuspended in PBS, and immediately analyzed by flow cytometry. Three biological replicates were performed in each experiment.

### Heat shock and killing assay

Cells at the exponential growth phase were harvested and subjected to heat shock by incubation at 42°C for 30 min. Immediately following heat treatment, the cultures were exposed to a lethal dose of meropenem (2 μg/ml) and incubated for 4 hours at 37°C. Surviving cells were quantified by serial dilution and plating on LB agar plates, followed by colony enumeration after overnight incubation. The killing assay after heat shock was performed with three biological replicates.

### Construction of *K. pneumoniae rpoS* gene deletion mutant

The *rpoS* gene knockout strain was generated via homologous recombination using the temperature-sensitive plasmid pKOV^44^. Genomic DNA from overnight cultures of the MGH66 WT strain was extracted using the Quick-DNA HMW MagBead Kit. DNA fragments (∼1 kb each) flanking the *rpoS* gene were amplified from MGH66 genomic DNA by PCR with Q5 High-Fidelity 2× Master Mix using primer pairs PMSJ087/PMSJ088 and PMSJ089/PMSJ090 (Supplementary Table 18). The pKOV plasmid backbone was similarly amplified using primers PMSJ064 and PMSJ065 (Supplementary Table 18). The three PCR products were assembled via NEBuilder HiFi DNA Assembly based on designed overlapping regions to generate the recombinant plasmid pKOV_up_down_*rpoS*. The resulting construct was transformed into *E. coli* NEB 10-beta and propagated at 30°C on LB agar supplemented with 25 μg/ml chloramphenicol (Sigma-Aldrich, C0378). Plasmid integrity was verified through whole-plasmid sequencing.

The confirmed pKOV_up_down_*rpoS* plasmid was introduced into *K. pneumoniae* MGH66 WT via electroporation. Transformants were selected on LB agar supplemented with 25 μg/ml chloramphenicol at 37°C. Owing to the temperature-sensitive origin of replication in pKOV, plasmid integration into the chromosome, via single-crossover recombination at either the upstream or downstream homology arm, was induced by shifting the incubation temperature to 42°C. To facilitate resolution of the integrated plasmid and achieve double-crossover recombination, integrant strains were cultured in LB without antibiotics at 37°C. Cultures were then diluted and plated onto LB agar without antibiotic. Individual colonies were screened for chloramphenicol sensitivity to identify clones that had lost the plasmid backbone. PCR screening using primers PMSJ087/PMSJ090 was performed to confirm deletion of the *rpoS* gene. The final *rpoS* depletion mutant strain MGH66:Δ*rpoS* was validated by whole-genome sequencing.

### Nutrient shift for killing assay

To investigate how nutrient shifts affect antibiotic persistence in stationary-phase *K. pneumoniae*, overnight cultures of the MGH66 WT strain were prepared by inoculating glycerol stocks into 80 ml of MHB in 500 ml flasks and incubating at 37°C with shaking. The following day, 1 ml of the culture was set aside, and the remaining volume was centrifuged at 4,500× g for 15 min. The supernatant was filtered using a 0.22 μm PES membrane (Millipore Express PLUS, S2GPU02RE) to obtain spent medium. The reserved overnight culture was diluted 1:100 into the spent medium. Aliquots of 1 ml were dispensed into 1.5 ml tubes and individually supplemented with either no additive or one of the following nutrients: 20 mM of glucose (Sigma-Aldrich, G8270), 10 mM of ammonium sulfate (Sigma-Aldrich, A4915), 10 mM of glutamine (Sigma-Aldrich, 1294808), 1% of casamino acids (Sigma-Aldrich, 65072-00-6), 2 mM of magnesium sulfate (Sigma-Aldrich, M2643), 10 mM Ammonium iron (II) sulfate hexahydrate (Sigma-Aldrich, 09719, simplified as FeSO_4_). Cultures were then treated with meropenem (2 μg/ml) or ciprofloxacin (2.5 μg/ml) and incubated at 37°C with shaking. After 4 hours of antibiotic exposure, cells were harvested by centrifugation, resuspended in 100 μl PBS, serially diluted, and plated on LB agar plate. Surviving cells were quantified by colony counting following overnight incubation. All experiments were performed in triplicate.

### Thiourea reverses Fe²^+^-enhanced ciprofloxacin killing assay

Stationary-phase *K. pneumoniae* MGH66 cells were diluted 1:100 into 1 ml spent medium in 1.5 ml tubes and supplemented with one of the following conditions: no additive, ciprofloxacin alone (2.5 μg/ml), FeSO_4_ alone (10 mM), thiourea alone (200 mM, Sigma-Aldrich, T8656), FeSO_4_ plus ciprofloxacin, FeSO_4_ plus thiourea, or FeSO_4_ with both ciprofloxacin and thiourea. Cultures were incubated at 37°C with shaking for 4 hours, then harvested by centrifugation, washed, serially diluted in PBS, and plated on LB agar. CFUs were enumerated after overnight incubation to quantify surviving cells. All conditions were performed in triplicate.

### Bulk RNA-seq

Stationary-phase *K. pneumoniae* MGH66 cells were diluted 1:100 into 1 ml of spent medium in 1.5 ml tubes and supplemented with either no additive, FeSO_4_ alone (10 mM), or FeSO_4_ combined with ciprofloxacin (2.5 μg/ml). Cultures were incubated at 37°C with shaking for 4 hours, after which cells were harvested by centrifugation and resuspended in 500 μl TRIzol (Invitrogen, 15596026) for total RNA extraction. RNA was purified using the Direct-zol RNA Miniprep Kit (Zymo, R2052) according to the supplied protocol. For library preparation, 100 ng of total RNA was used with the NEBNext Ultra II Directional RNA Library Prep Kit (NEB, E7760S). Sequencing was performed in paired-end mode on an Illumina NextSeq2000 platform.

## Computational analysis

### Processing of raw sequencing data

Raw sequencing reads were trimmed and quality-checked using *fastp* (v0.19.6)^45^. Demultiplexing was performed based on CB1 with a custom shell script to separate samples. Cell barcodes and UMI sequences were extracted from the reads and incorporated into read headers using *UMI-tools* (v1.1.4)^46^. Processed reads were then aligned to the *Klebsiella pneumoniae* MGH 66 strain genome (BioProject: PRJNA1309751) using *BWA* (v0.7.17)^47^. Gene-level quantification was performed with *featureCounts* (v2.0.5)^48^, and the final count table was generated with *UMI-tools*.

### Analysis of BacDrop_v2 data

For each scRNA-seq experiment using BacDrop_V2, approximately 100,000 cells were barcoded per sample, and sequencing was performed at a depth of ∼2,000–3,000 reads per cell. For downstream analysis, the top 10,000 cells with the highest UMI counts were selected for each sample, consistent with the published BacDrop data analysis pipeline^23^, which prioritizes cells with sufficient transcript coverage for robust state identification in bacterial scRNA-seq. Count tables were analyzed using the *Seurat* package (v5.1.0)^49^. Quality control was performed by filtering out low-quality cells with fewer than 15 detected genes, as well as potential barcode collisions represented by cells with more than 300 detected genes. Gene counts were normalized using default *Seurat* parameters, and the top 2,000 variable features were identified with the variance-stabilizing transformation (vst) method. All genes were then scaled across cells, and dimensionality reduction and clustering were performed by principal component analysis (PCA) on the variable features, followed by UMAP. Marker genes for each cluster were identified with *FindAllMarkers* using a log_2_ fold change threshold of 0.5. Cell states were annotated by GO and pathway enrichment analysis of cluster marker genes (log_2_ fold change > 1, adjusted p < 0.05) using STRING (v12.0)^50^.

### Bulk RNA-seq analysis

To assess potential bias during single-cell RNA library construction, we performed correlation analysis between bulk RNA-seq and scRNA-seq data. Cell barcode and UMI sequences were removed from scRNA-seq reads using *fastp* (v0.24.0). These trimmed reads, together with bulk RNA-seq reads, were aligned to the reference genome with *BWA* (v0.7.19). Read counts were quantified using *featureCounts* (v2.1.1) to generate a raw gene count matrix, and gene-level correlations were visualized with *Matplotlib Pyplot*.

For bulk RNA-seq differential expression analysis, raw sequencing reads were quality-checked with *fastp* (v0.24.0) and aligned to the reference genome using *BWA* (v0.7.19). Gene-level expression matrices were generated with *featureCounts* (v2.1.1), and raw counts were normalized with the *rlog* function in the *DESeq2* package (v1.46.0)^51^. Normalized expression data were visualized with the *ComplexHeatmap* package. Differentially expressed genes were identified using *DESeq2*, with adjusted p values and fold changes calculated. Volcano plots were generated with *ggplot2*.

## Acknowledgements

We thank Drs. Elaine Tuomanen, Richard Lee, David Rogers, Jinna Bai for comments on the manuscript. We thank the Hartwell Center at St. Jude Children’s Research Hospital for sequencing, and the flow core for flow cytometry analysis. This research is funded by National Institute of General Medical Sciences grant R35GM157243 to P.M. and Children’s Infection Defense Center at St. Jude Children’s Research Hospital (PROJ-1001461) to P.M.

## Author contributions

Q.L., X.L. and P.M. conceived the study and performed the experiments. Q.L., Z.Y., A.I., T.S., J.W.R. and P.M. developed the methodology. Q.L., Z.Y. and P.M. performed data analysis and visualization. Q.L., Z.Y. and P.M. wrote the manuscript draft. Q.L., Z.Y., J.W.R. and P.M. revised and edited the manuscript. J.W.R. and P.M. supervised the study. P.M. provided the funding and resource.

## Materials & Correspondence

The bacterial strains and constructs will be available upon request to P.M. after the publication of this manuscript.

## Statistical information

All statistical analyses were carried out using GraphPad Prism 10. One-way ANOVA was applied to compare means across multiple groups, while unpaired *t*-tests were used for comparisons between two independent groups. Sample sizes, statistical tests, and measures of precision are reported in the figure legends and results sections.

## Data Availability

The processed counting matrix of all scRNA-seq experiments have been deposited to GEO repository (GEO: GSE306416). The raw sequencing data of bulk RNA-seq and scRNA-seq experiments have been deposited to SRA depository (PRJNA1310171). All data will be publicly available upon the publication of this manuscript. All scRNA and bulk RNA analysis scripts are deposited at https://github.com/Zehui-stjude/BacDrop_v2 and will be publicly available upon the publication of this manuscript.

