## Supplemental figures 1 to 17 for "Antibiotic Persistence Emerges from Cell-State-Driven Transcriptional Reprogramming"

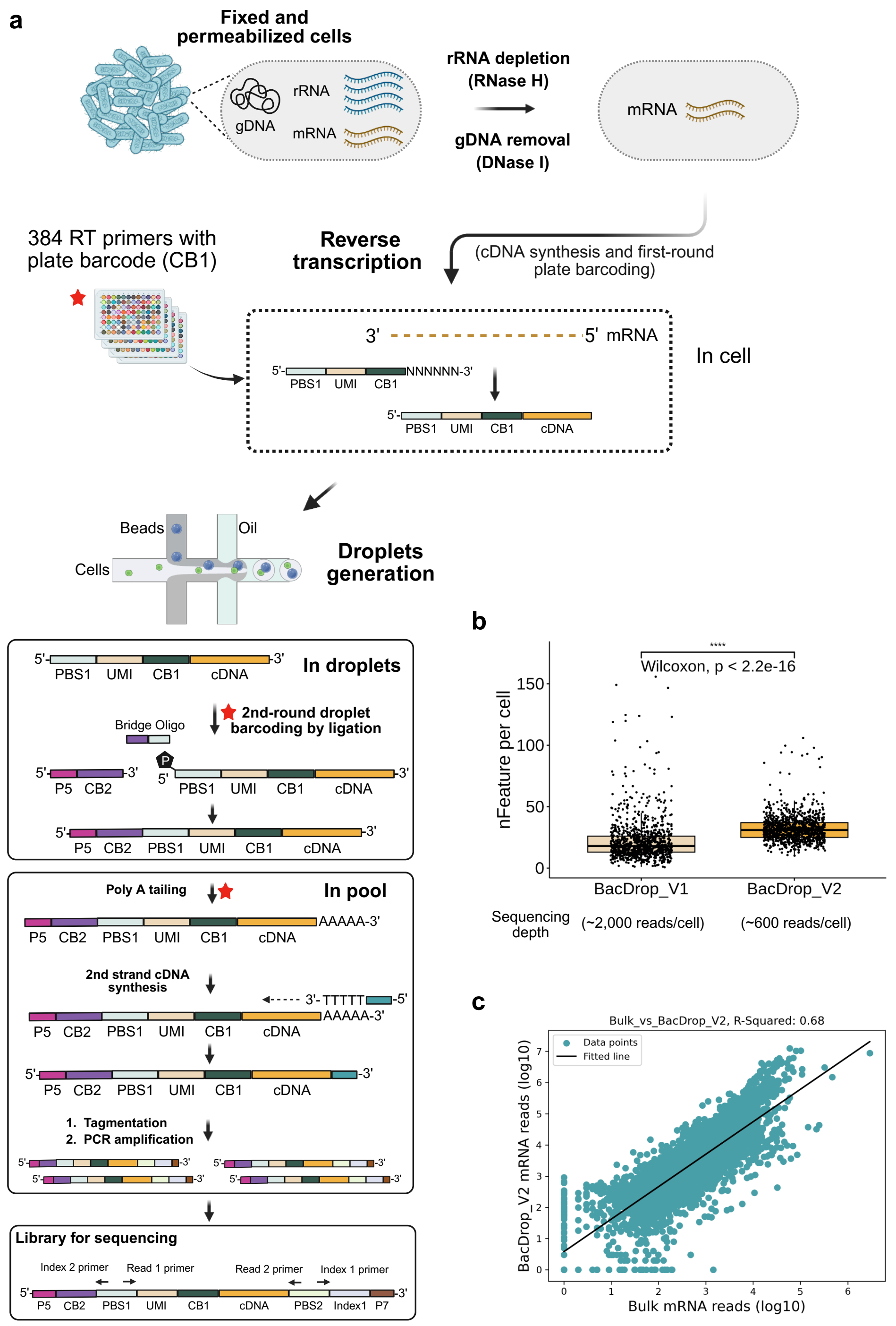


**Supplementary Figure 1. Workflow and performance of the modified BacDrop method (BacDrop_V2). a**, Overview of the BacDrop_V2 workflow and red asterisks indicate the steps that differ from BacDrop_V1. Following fixation and permeabilization, rRNA and genomic DNA is depleted *in situ* using RNase H and DNase I treatment, respectively. In-cell reverse transcription (RT) is performed in 96-well plates using primers comprising a phosphate group at the 5', a partial Illumina primer-binding sequence (PBS1), an 8-bp unique molecular identifier (UMI), a 13-bp first-round cell barcode (CB1), and a 6-bp random sequence for RT priming. This step generates cDNA and introduces CB1 in-cell. After RT, cells are pooled and encapsulated with10x gel beads from the ATAC-seq kit for round-2 droplet barcoding. Round-2 droplet barcoding is achieved by thermo-ligation of a second barcode (CB2) to the 5′ end of cDNA via a bridge oligo co-encapsulated in droplets, generating molecules that carry CB1–UMI–cDNA together with CB2. The barcoded cDNA is then recovered from droplets, pooled, purified, and poly(A)-tailed at the 3′ end, enabling second-strand synthesis using the SMRT_dT primer. Double-stranded cDNA molecules containing CB1, CB2 and UMI, flanked by adaptor sequences, are purified and amplified. Final library construction is completed by tagmentation and PCR enrichment to generate Illumina-compatible sequencing libraries. Each cell is uniquely identified by the combinatorial CB1 plus CB2 barcode pair. Red asterisks indicate the steps that differ from BacDrop_V1, specifically: (i) omission of the in-cell poly(A)-tailing step prior to droplet encapsulation; (ii) replacement of the BacDrop_V1 polymerase-mediated CB2 writing step during oligo(dT)-primed second-strand synthesis with a ligation-based in-droplet CB2 assignment mediated by a bridge oligo; (iii) poly(A)-tailing after the cDNA, now barcoded with both CB1 and CB2, was pooled and purified. **b**, Comparison of mRNA features detected per cell between BacDrop_V1 and BacDrop_V2. Approximately 1,000 cells were analyzed for each method. BacDrop_V1 was sequenced at an average depth of ~2,000 reads per cell and yielded a mean of 22 mRNA genes detected per cell, whereas BacDrop_V2 yielded a mean of 32 mRNA genes detected per cell at ~600 reads per cell, indicating improved sensitivity per sequencing depth in BacDrop_V2. (Center lines indicate medians; box bounds indicate the interquartile range; whiskers indicate the full range unless otherwise noted.) **c**, Concordance between BacDrop_V2 and matched bulk RNA-seq for *K. pneumoniae* MGH66, shown as log10-transformed mRNA read counts ($R^2 = 0.68$).

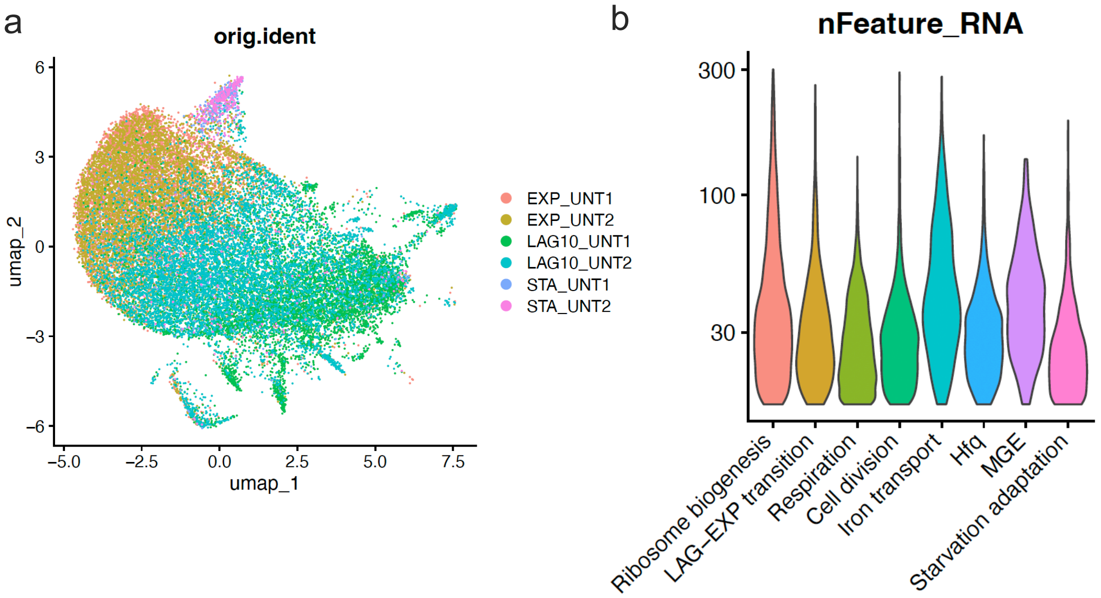


**Supplementary Figure 2. UMAP reproducibility and mRNA content across cell states in untreated samples. a**, UMAP projection of single-cell RNA-seq data from *K. pneumoniae* MGH66, colored by sample identity. Samples correspond to cells collected from exponential (EXP), early lag (LAG10), and stationary (STA) phases under untreated (UNT) conditions. Two biological replicates were included for each condition. **b**, Violin plots showing the distribution of detected genes (excluding rRNA and tRNA; nFeature_RNA) per cell in selected functional groups, including ribosome biogenesis, LAG-EXP transition, respiration, cell division, iron transport, a small RNA chaperone (Hfq), mobile genetic elements (MGE), and starvation adaptation.

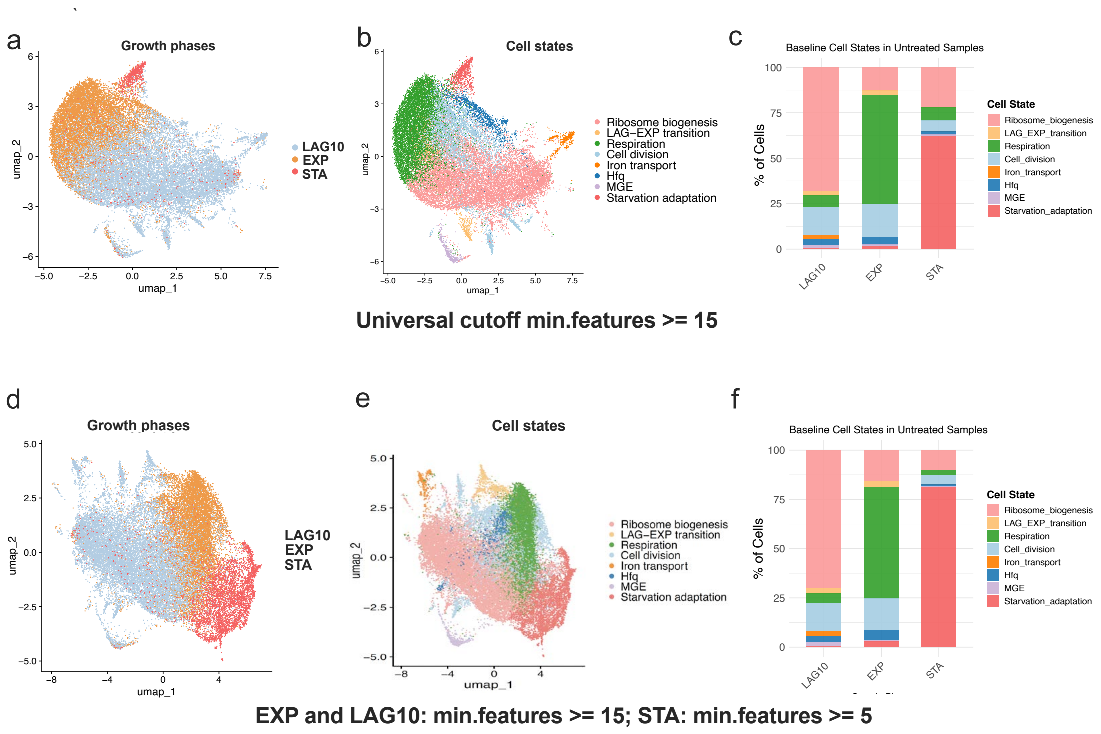


**Supplementary Figure 3. Effect of minimal feature thresholds on scRNA-seq state identification in untreated samples.** scRNA-seq analysis of untreated *Klebsiella pneumoniae* MGH66 cells under different minimal feature cutoffs for cell inclusion. **a - c**, Analyses using a universal cutoff of min.features ≥ 15 for all growth phases. **a**, UMAP colored by growth phase: lag (LAG10), exponential (EXP), and stationary (STA). **b**, The same UMAP colored by transcriptional cell state. **c**, Relative abundance of each transcriptional state within each growth phase, shown as the percentage of cells. **d - f**, Analyses using phase-specific cutoffs, with min.features ≥ 15 for EXP and LAG10 cells and min.features ≥ 5 for STA cells. **d**, UMAP colored by growth phase. **e**, UMAP colored by transcriptional cell state. **f**, Relative abundance of each transcriptional states within each growth phase. All conditions include two biological replicates; bar plots show the average across replicates.

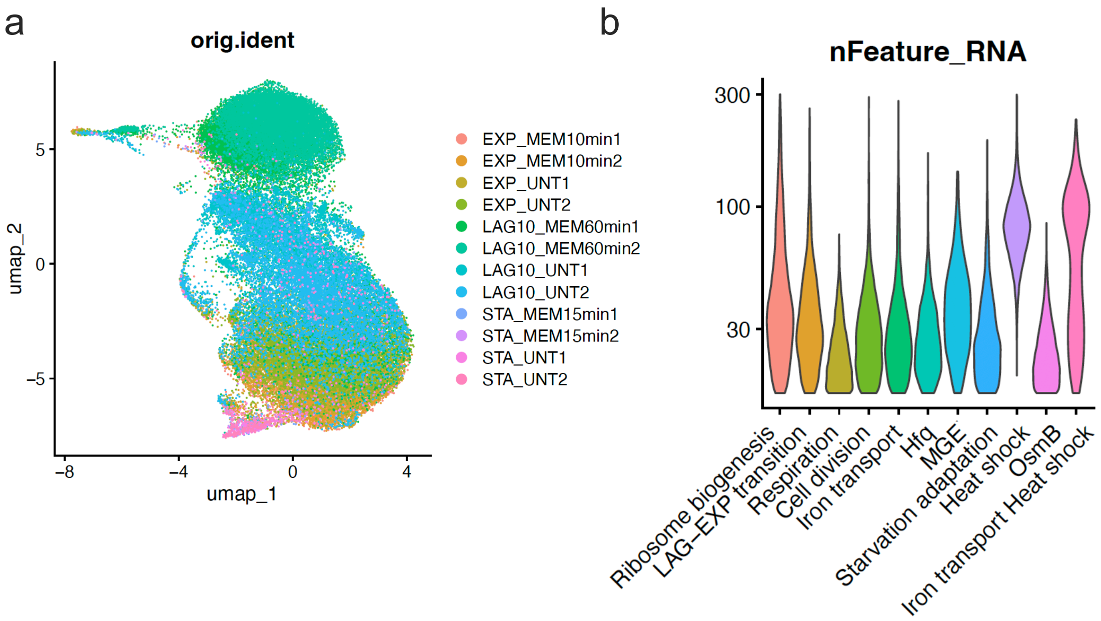


**Supplementary Figure 4. UMAP reproducibility and mRNA content across cell states in untreated and meropenem-treated samples. a,** UMAP visualization of single-cell transcriptomes from *K. pneumoniae* MGH66 collected during exponential (EXP), early lag (LAG10), and stationary (STA) phases under untreated (UNT) or meropenem-treated (MEM) conditions (2 μg/ml) for the indicated durations (10, 60, or 15 min). Points represent individual cells, colored by sample identity. Two biological replicates were included for each condition. **b,** Violin plots showing the distribution of detected genes (excluding rRNA and tRNA; nFeature_RNA) per cell within functional categories, including ribosome biogenesis, LAG-EXP transition, respiration, cell division, iron transport, Hfq, MGE, starvation adaptation, OsmB, and iron transport combined with heat shock.

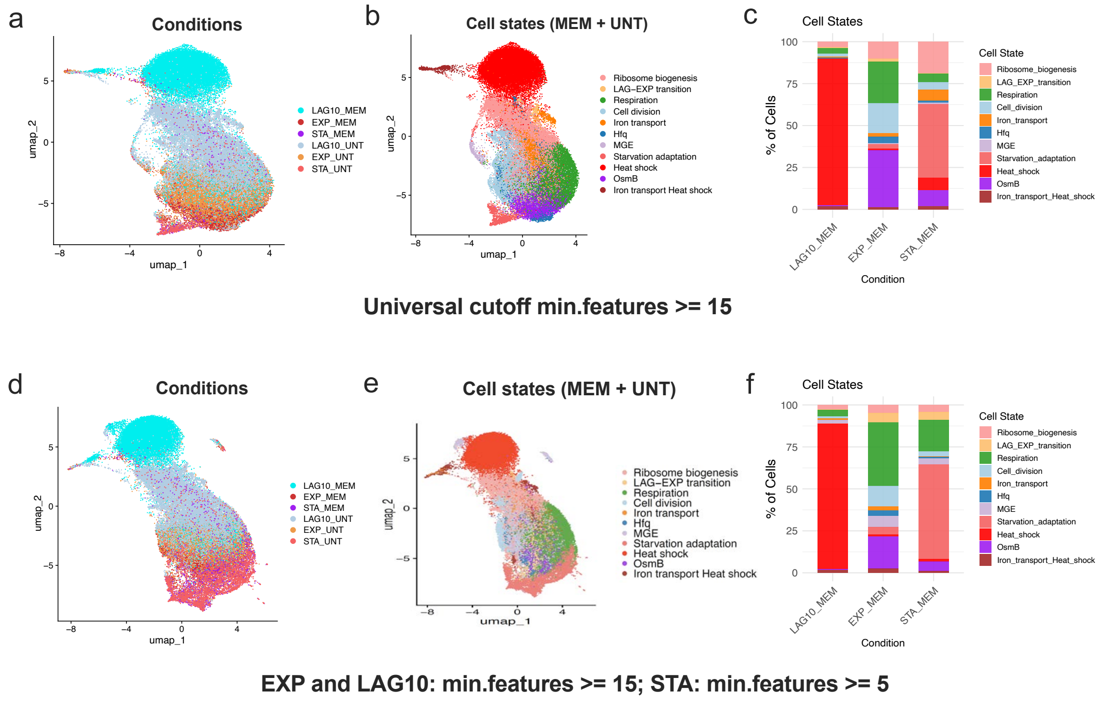


**Supplementary Figure 5. Effect of minimal feature thresholds on scRNA-seq state identification in meropenem-treated samples.** scRNA-seq analysis of *Klebsiella pneumoniae* MGH66 cells treated with meropenem (MEM) together with untreated controls (UNT) under different minimal feature cutoffs for cell inclusion. **a - c**, Analyses using a universal cutoff of min.features ≥ 15 for all growth phases. **a**, UMAP colored by condition (LAG10_MEM, EXP_MEM, STA_MEM, LAG10_UNT, EXP_UNT, STA_UNT). **b**, The same UMAP colored by transcriptional cell state. **c**, Relative abundance of each transcriptional state within each meropenem-treated condition, shown as the percentage of cells. **d - f**, Analyses using phase-specific cutoffs, with min.features ≥ 15 for EXP and LAG10 cells and min.features ≥ 5 for STA cells. **d**, UMAP colored by condition. **e**, UMAP colored by transcriptional cell state. **f**, Relative abundance of each transcriptional state within each meropenem-treated condition. All conditions include two biological replicates; bar plots show the average across replicates.

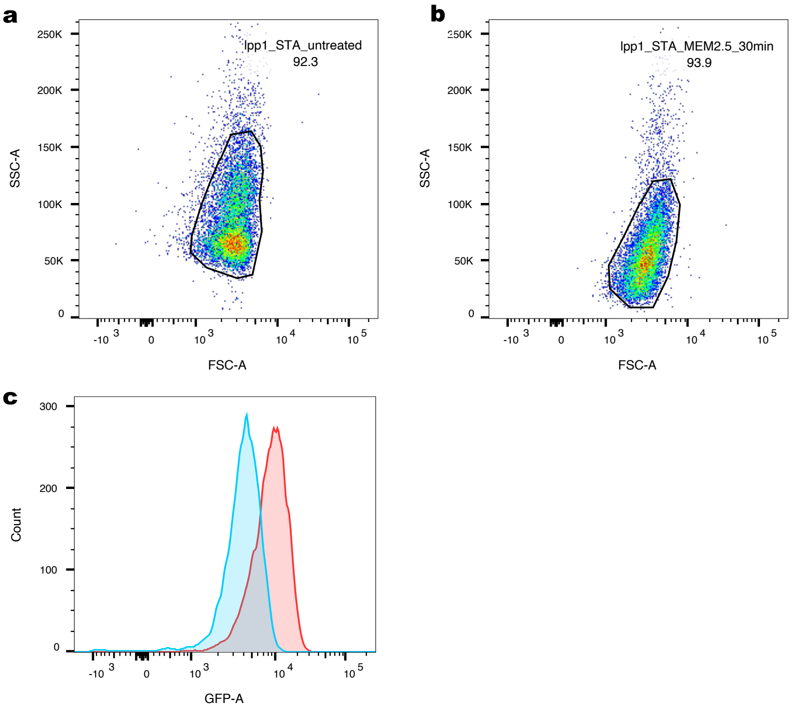


| | Sample Name | Subset Name | Count |
|---|---|---|---|
| (blue) | 941680.fcs | lpp1_STA_untreated | 9046 |
| (red) | 942704.fcs | lpp1_STA_MEM2.5_30min | 9356 |

**Supplementary Figure 6. Flow cytometry analysis of *lpp1* induction in stationary-phase (STA) cells following meropenem treatment. a-b,** Gating of *K. pneumoniae* MGH66 P*lpp1:gfp* stationary-phase cells without treatment (a) or after exposure to meropenem (2 µg/ml) for 30 min (b). **c**, Overlay of fluorescence intensity histograms from untreated (blue) and CIP-treated (pink) STA cells, confirming a rightward shift in the meropenem-treated population indicative of *lpp1* induction.

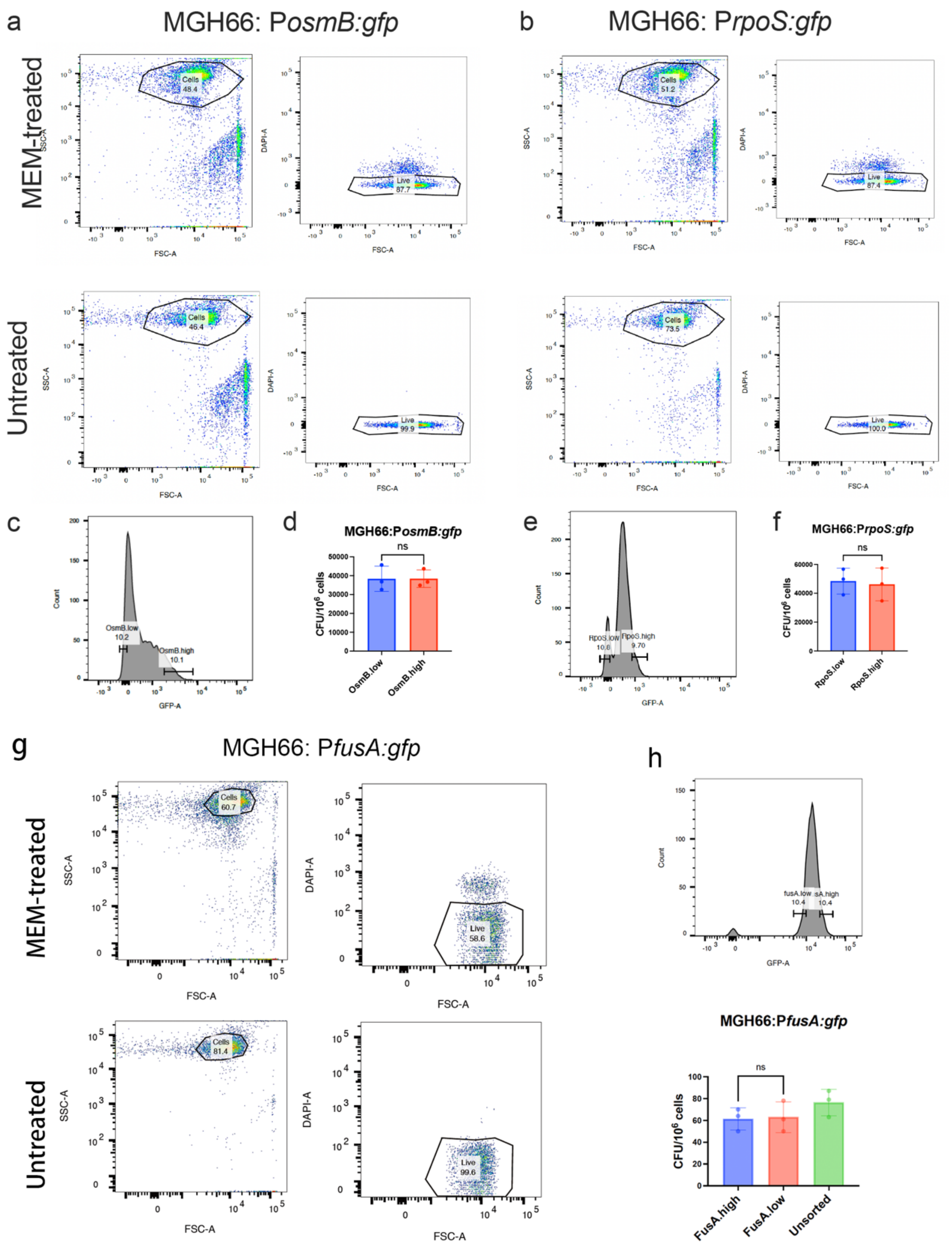


**Supplementary Figure 7. Flow cytometry gating to validate heterogeneity in meropenem-treated exponential-phase (EXP) cells. a, b, g**, *K. pneumoniae* MGH66 WT strain carrying P*osmB:gfp* (a), P*rpoS:gfp* (b), or P*fusA:gfp* (g) were analyzed by flow cytometry under meropenem (MEM)-treated (2 µg/ml) and untreated conditions. Left plots show gating of total cell populations based on forward scatter (FSC-A) and side scatter (SSC-A) to exclude cell debris; right plots show gating of DAPI-negative events to select live cells. Percentages indicate the proportion of events within each gate relative to the parent population. **c-f,** Sorted cells showed comparable survival after sorting into MHB medium without antibiotics. In parallel with sorting cells into MEM-supplemented MHB medium to assess survival under meropenem treatment (2 µg/ml) (Fig. 2g), untreated controls were included to confirm that sorting itself did not affect viability. GFP-low and GFP-high subpopulations from MEM-treated *MGH66*: P*osmB:gfp* (c, d) or MGH66: P*rpoS:gfp* cultures (e, f) were gated for survival assays. A total of 10^6 live cells from each subpopulation were sorted into 1 ml LB medium without antibiotics and immediately plated on LB agar. Colony-forming units (CFUs) were enumerated after 24 h of incubation. All assays were performed in triplicate (Welch's *t*-test, *n* = 3). Mean ± s.e.m. is shown. **h,** A positive control GFP reporter strain MGH66:P*fusA:gfp* was treated with meropenem (2 ug/ml) and the survival rates of FusA.low, FusA.high, and unsorted cells were compared under meropenem treatment (2 µg/ml). No significant difference on their survival rates was observed between the FusA.low and FusA.high subpopulations (Welch's t-test, n = 3). These data indicate that differential survival observed for stress-responsive reporters is not attributable to nonspecific GFP accumulation.

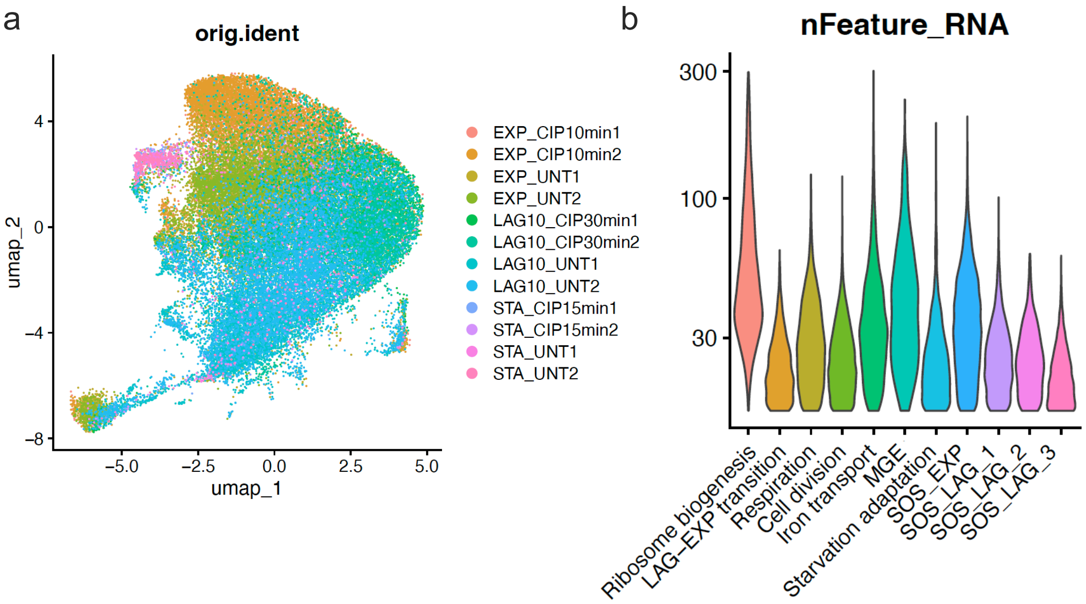


**Supplementary Figure 8. UMAP reproducibility and mRNA content across cell states in untreated and ciprofloxacin-treated samples.** **a**, UMAP visualization of single-cell transcriptomes from *K. pneumoniae* MGH66 collected during exponential (EXP), early lag (LAG10), and stationary (STA) phases under untreated (UNT) or ciprofloxacin-treated (CIP) (2.5 μg/ml) conditions for the indicated durations (10, 30, or 15 min). Points represent individual cells, colored by sample identity. Two biological replicates were included for each condition. **b**, Violin plots showing the distribution of detected genes (excluding rRNA and tRNA; nFeature_RNA) per cell within functional categories, including ribosome biogenesis, LAG-EXP transition, respiration, cell division, iron transport, MGE, starvation adaptation, SOS from EXP (SOS_EXP) and LAG (SOS_LAG_1 to 3).

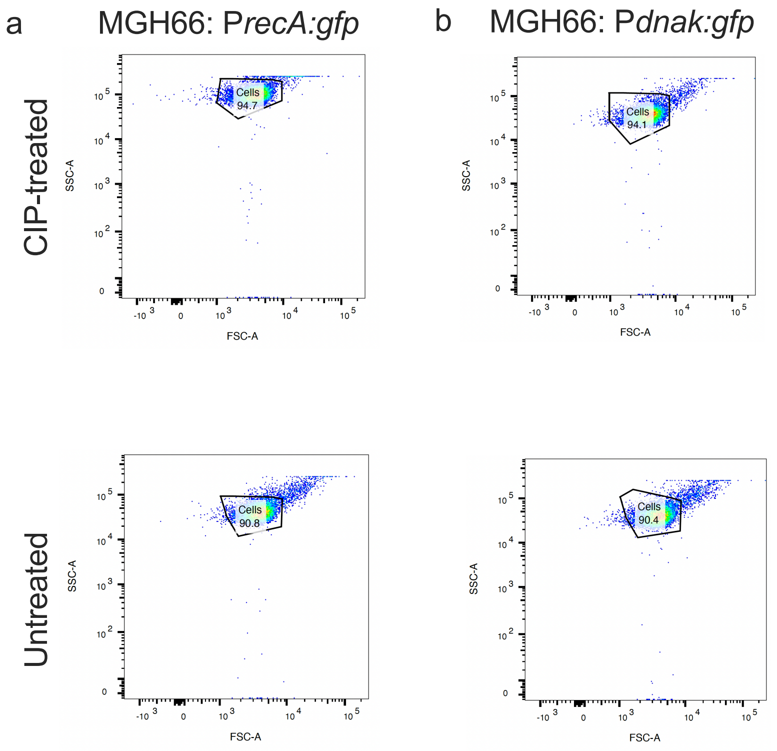


**Supplementary Figure 9. Flow cytometry gating to validate co-expression of *recA* and *dnaK* in ciprofloxacin-treated EXP cells. a,b**, *K. pneumoniae* MGH66 P*recA:gfp* and MGH66 P*dnak:gfp* strains were analyzed under ciprofloxacin-treated (2.5 µg/ml) (top panels) or untreated (bottom panels) conditions. Forward scatter area (FSC-A, x-axis) and side scatter area (SSC-A, y-axis) were used to distinguish bacterial cell populations from debris. The gated region represents intact single cells, defined by characteristic FSC-A/SSC-A profiles. Numbers indicate the percentage of events within each gate relative to the total detected events. DAPI staining was not included because ciprofloxacin-treated cells at this sampling time resisted to DAPI staining, limiting DAPI from efficiently entering the cells, although ~50% of cells were confirmed dead by plating.

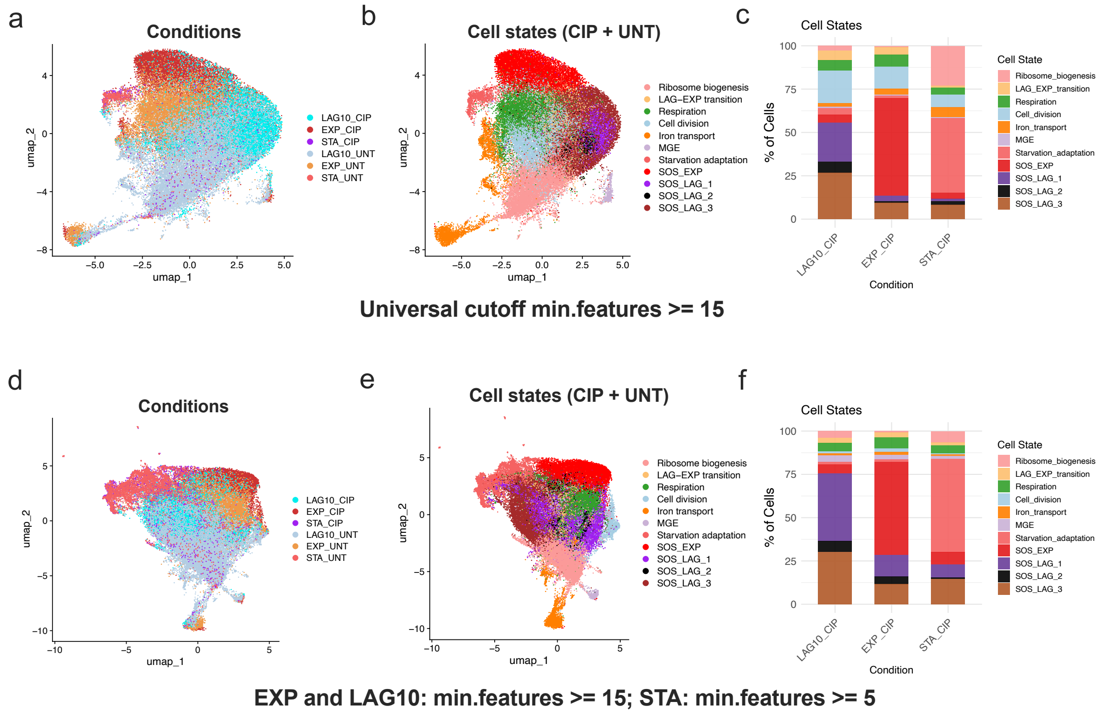


**Supplementary Figure 10. Effect of minimal feature thresholds on scRNA-seq state identification in ciprofloxacin-treated samples.** scRNA-seq analysis of *Klebsiella pneumoniae* MGH66 cells treated with ciprofloxacin (CIP) together with untreated controls (UNT) under different minimal feature cutoffs for cell inclusion. **a - c**, Analyses using a universal cutoff of min.features ≥ 15 for all growth phases. **a**, UMAP colored by condition (LAG10_CIP, EXP_CIP, STA_CIP, LAG10_UNT, EXP_UNT, STA_UNT). **b**, The same UMAP colored by transcriptional cell state. **c**, Relative abundance of each transcriptional state within each ciprofloxacin-treated condition, shown as the percentage of cells. **d - f**, Analyses using phase-specific cutoffs, with min.features ≥ 15 for EXP and LAG10 cells and min.features ≥ 5 for STA cells. **d**, UMAP colored by condition. **e**, UMAP colored by transcriptional cell state. **f**, Relative abundance of each transcriptional state within each ciprofloxacin-treated condition. All conditions include two biological replicates; bar plots show the average across replicates.

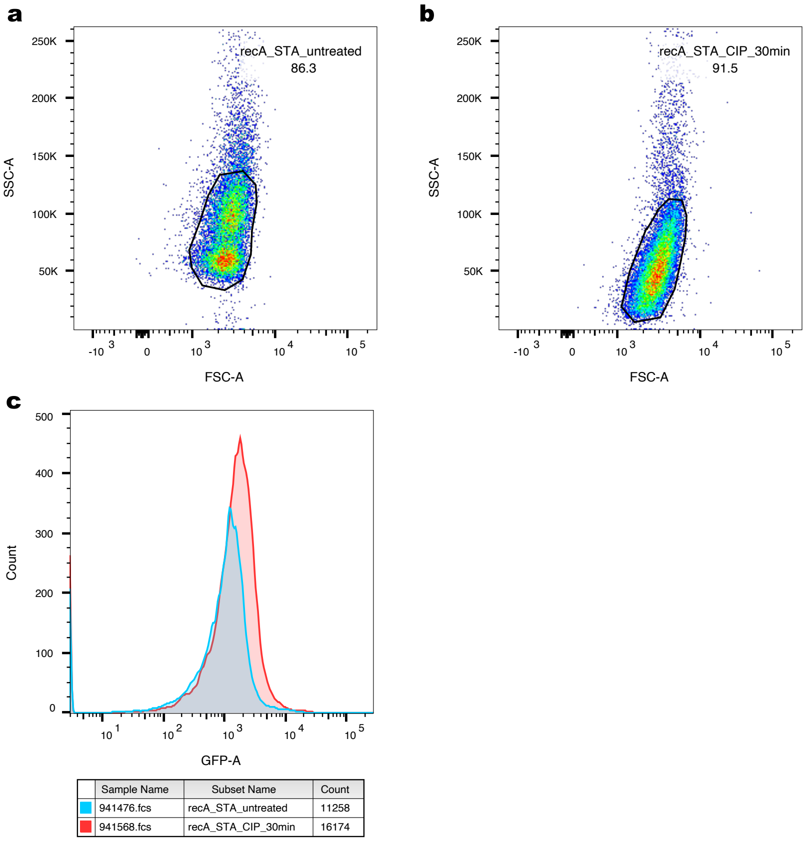


**Supplementary Figure 11. Flow cytometry analysis of *recA* induction in stationary-phase cells following ciprofloxacin treatment. a,b,** Gating of *K. pneumoniae* MGH66:P*recA:gfp* stationary-phase (STA) cells without treatment (a) or after exposure to ciprofloxacin (CIP) (2.5 µg $ml^{-1}$) for 30 min (b). **c**, Overlay of fluorescence intensity histograms from untreated (blue) and CIP-treated (pink) STA cells, confirming a rightward shift in the ciprofloxacin-treated population indicative of *recA* induction.

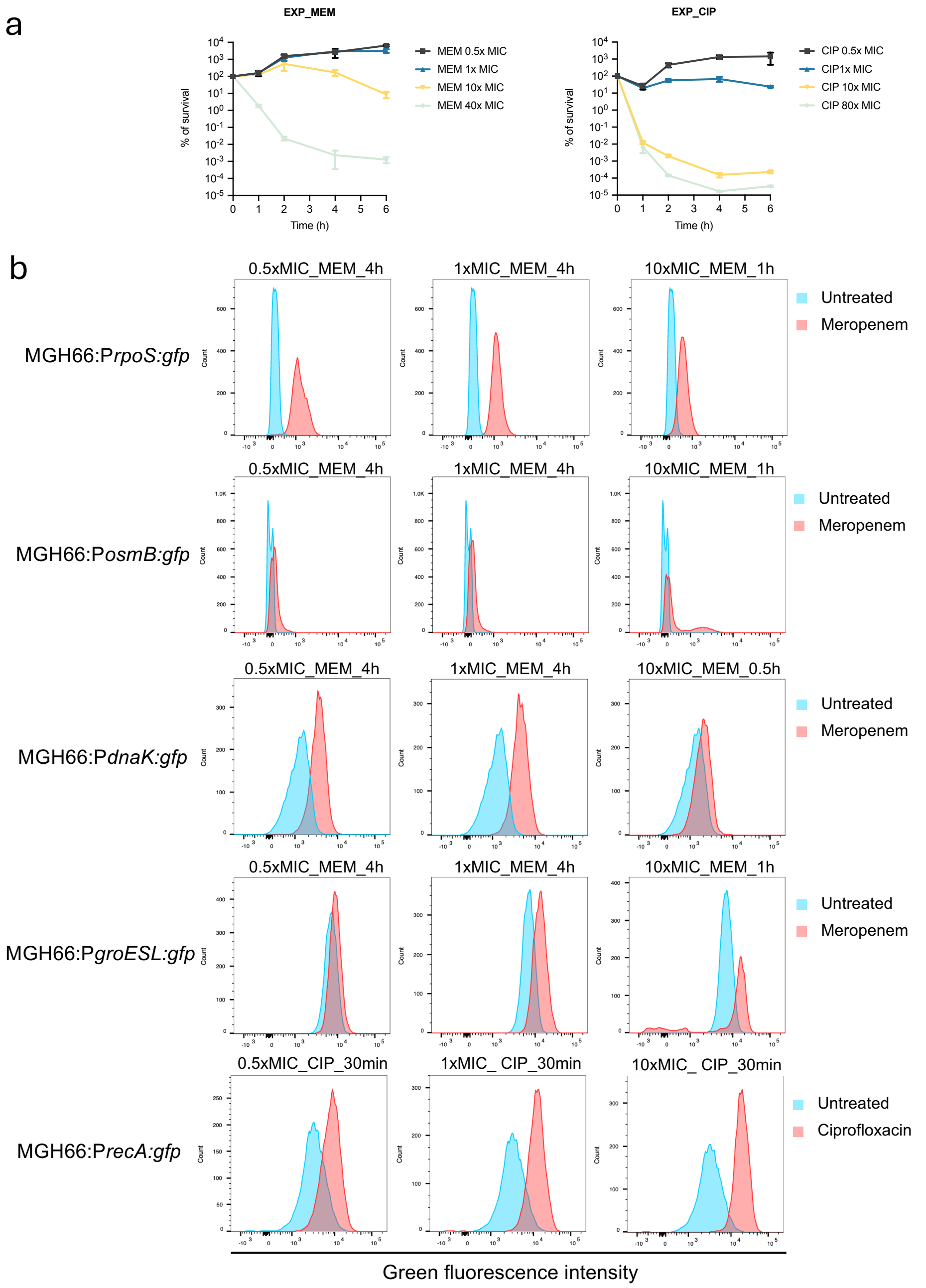


**Supplementary Figure 12. Killing dynamics of *K. pneumoniae* MGH66 under various antibiotic concentrations and corresponding flow-cytometry analysis of different reporter strains under these conditions. a.** Time-course survival curves of exponential-phase (EXP) *K. pneumoniae* MGH66 treated with 0.5×, 1×, 10×, or 40× the minimum inhibitory concentration (MIC) of meropenem (MEM; left) or 0.5×, 1×, 10×, or 80× MIC of ciprofloxacin (CIP; right). Survival is plotted as the percentage of the initial population remaining viable over time. **b.** Overlay histograms of GFP fluorescence measured by flow cytometry for GFP reporter strains

(MGH66:P*rpoS:gfp*, MGH66:P*osmB:gfp*, MGH66:P*dnaK:gfp*, MGH66:P*groESL:gfp*, and MGH66:P*recA:gfp*), following treatment with MEM or CIP at the indicated multiples of the MIC and time points shown above each panel. Untreated controls are shown in blue, and antibiotic-treated populations are shown in red. The x-axis indicates green fluorescence intensity.

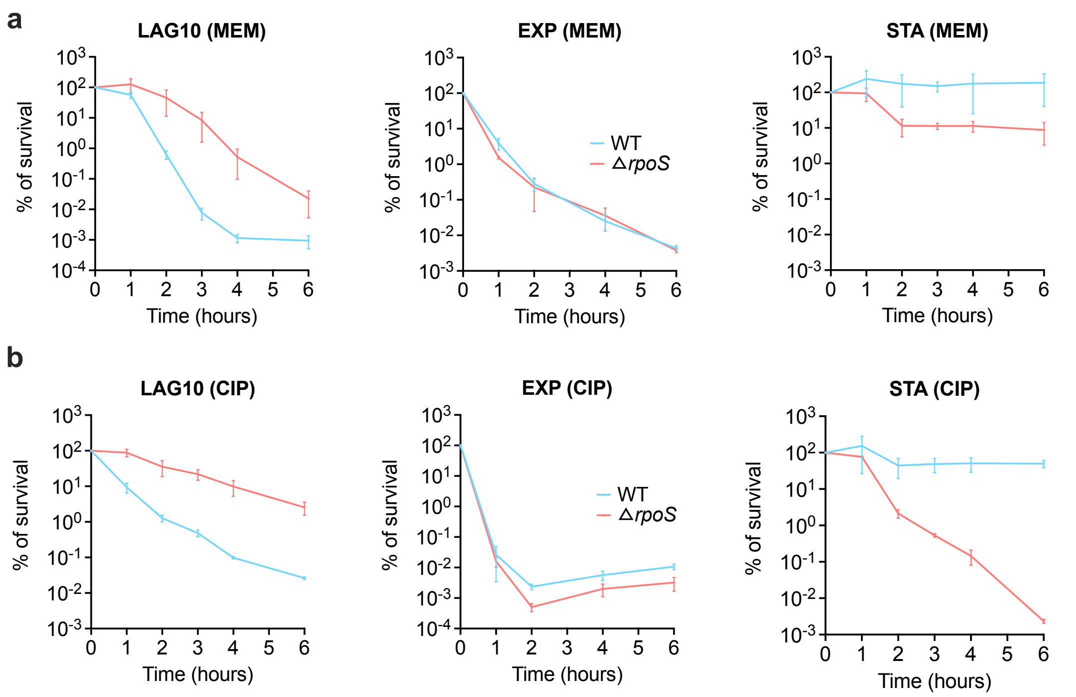


**Supplementary Figure 13. Killing dynamics of the wild-type (WT) *E. coli* K-12 strain and rpoS deletion strain (Δ*rpoS*) under meropenem (MEM) or ciprofloxacin (CIP) treatment. a,b**, Survival kinetics of *E. coli* K-12 WT (blue) and Δ*rpoS* (red) strains after exposure to (a) meropenem (2 μg/ml, 40× MIC; top) or (b) ciprofloxacin (2.5 μg/ml, 80× MIC; bottom) during exponential (EXP), early lag (LAG10), and stationary (STA) phases. Deletion of *rpoS* led to reduced level of persistence in the STA phase, elevated persistence in the LAG10 phase, and had minimal impact during the EXP phase. Data represent the mean of three independent biological replicates (n = 3), with error bars indicating standard deviations.

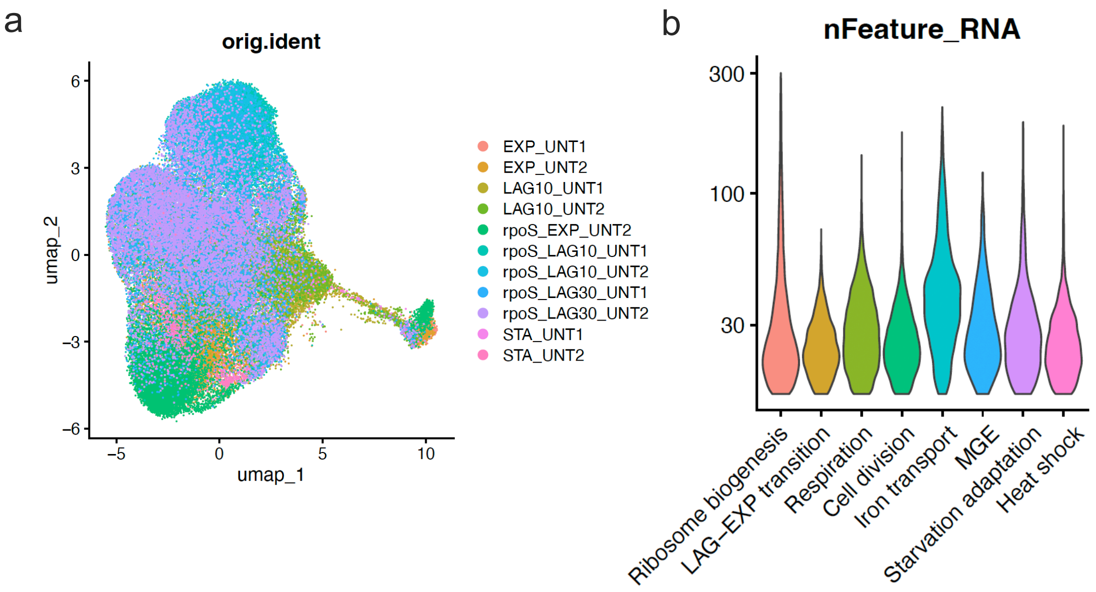


**Supplementary Figure 14. UMAP reproducibility and mRNA content across cell states in untreated samples of the WT and Δ*rpoS* strains. a**, UMAP visualization of single-cell transcriptomes from *K. pneumoniae* MGH66 WT or Δ*rpoS* strains collected during exponential (EXP), early lag (LAG10), and stationary (STA) phases under untreated (UNT) conditions. Points represent individual cells, colored by sample identity. Two biological replicates were included for each condition. **b**, Violin plots showing the distribution of detected genes (excluding rRNA and tRNA; nFeature_RNA) per cell within functional categories, including ribosome biogenesis, LAG-EXP transition, respiration, cell division, iron transport, MGE, starvation adaptation, and heat shock.

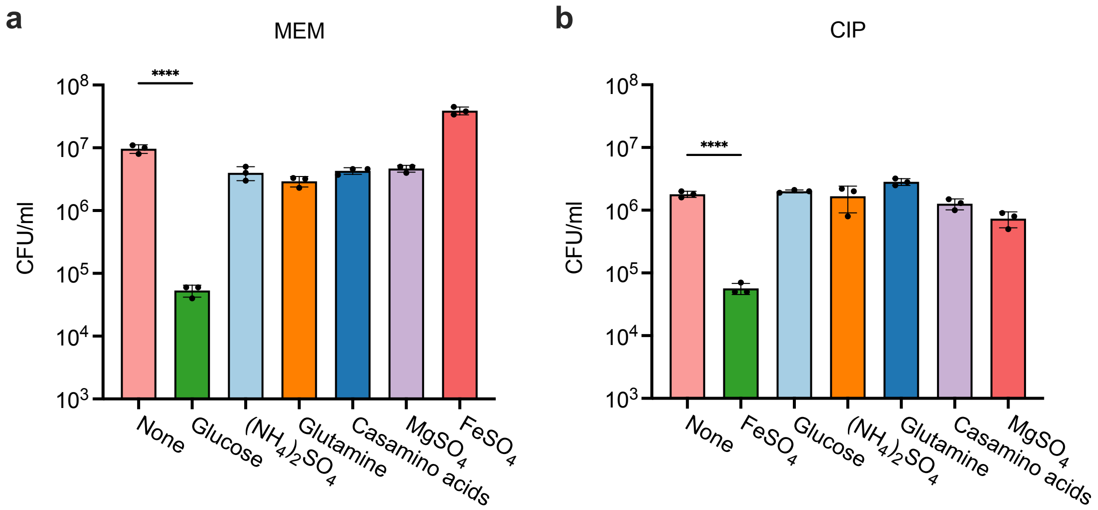


**Supplementary Figure 15. Survival of stationary-phase (STA) cells under meropenem (MEM) and ciprofloxacin (CIP) treatment with various nutrient supplements. a,b**, STA cells from overnight cultures were diluted into either spent medium alone or spent medium supplemented individually with glucose (20 mM), $(NH_4)_2SO_4$ (10 mM), glutamine (10 mM), casamino acids (1%), $MgSO_4$ (2 mM), or $FeSO_4$ (10 mM), followed by treatment with meropenem (2 µg/ml) (a) or ciprofloxacin (2.5 µg/ml) (b). CFUs were determined by plating cultures on LB agar after 4 hours of antibiotic exposure. All assays were performed in triplicate. (*****p* < 0.0001, lognormal Welch's t test, n=3). Mean and standard error are plotted.

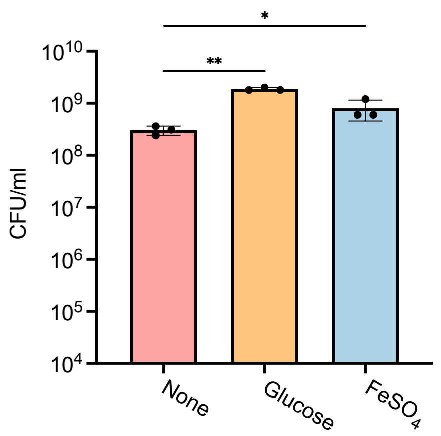


**Supplementary Figure 16. Glucose or $FeSO_4$ alone did not impact survival of *K. pneumoniae* MGH66 WT in the absence of antibiotics.** Staionary-phase (STA) cells from overnight cultures were diluted into spent medium supplemented with no additive, glucose (20 mM) or $FeSO_4$ (10 mM), and incubated for 4 hours. CFUs were determined by plating cultures on LB agar. All experiments were performed in triplicate. (*$p < 0.05$, **$p < 0.01$, lognormal Welch's t test, n=3). Mean and standard error shown.

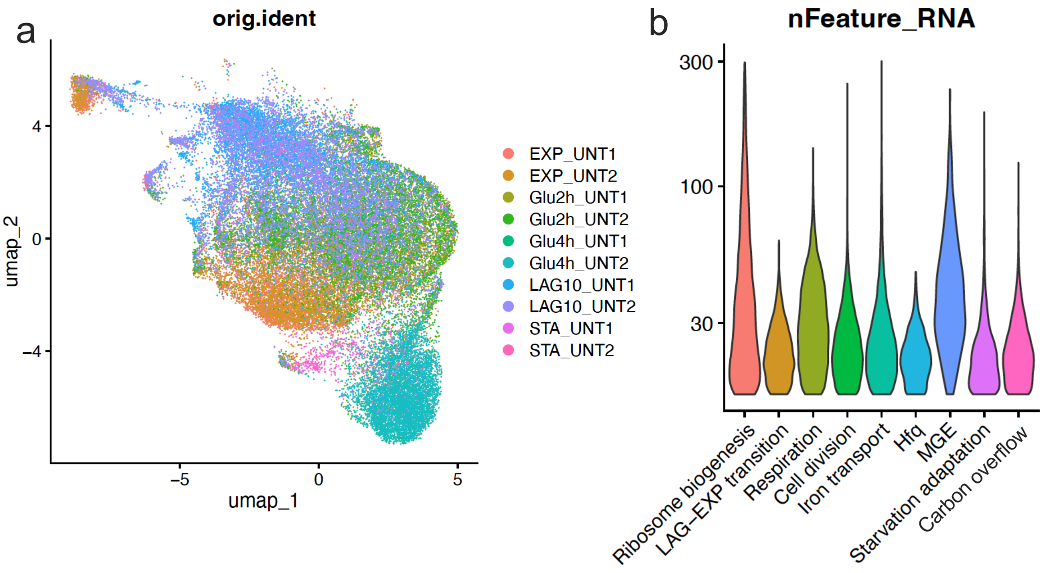


**Supplementary Figure 17. UMAP reproducibility and mRNA content across cell states in untreated samples across growth phases and stationary-phase (STA) cells with glucose supplementation. a**, UMAP visualization of single-cell transcriptomes from *K. pneumoniae* MGH66 WT collected during exponential (EXP), early lag (LAG10), and stationary (STA) phases, as well as STA cells grown with 20 mM glucose for 2 hours (Glu2h) or 4 hours (Glu4h). Points represent individual cells, colored by sample identity. Two biological replicates were included for each condition. **b**, Violin plots showing the distribution of detected genes (excluding rRNA and tRNA; nFeature_RNA) per cell within functional categories, including ribosome biogenesis, LAG-EXP transition, respiration, cell division, iron transport, Hfq, MGE, starvation adaptation, and carbon overflow.
